# The actin-modulating protein Synaptopodin mediates long-term survival of dendritic spines

**DOI:** 10.1101/2020.05.08.080374

**Authors:** Kenrick Yap, Alexander Drakew, Dinko Smilovic, Michael Rietsche, Mario Vuksic, Domenico Del Turco, Thomas Deller

## Abstract

Large spines are stable and important for memory trace formation. The majority of large spines also contains Synaptopodin (SP), an actin-modulating and plasticity-related protein. Since SP stabilizes F-actin, we speculated that the presence of SP within large spines could explain their long lifetime. Indeed, using time-lapse 2-photon-imaging of SP-transgenic granule cells in mouse organotypic tissue cultures we found that spines containing SP survived considerably longer than spines of equal size without SP. Of note, SP-positive spines that underwent pruning first lost SP before disappearing. Whereas the survival time courses of SP-positive (SP+) spines followed conditional two-phase decay functions, SP-negative (SP-) spines and all spines of SP-deficient animals showed single exponential decays. These results implicate SP as a major regulator of long-term spine stability: SP clusters stabilize spines and the presence of SP indicates spines of high stability.

## Introduction

Dendritic spines are protrusions found on the majority of excitatory neurons in vertebrate brains. They form characteristic axo-spinous synapses and play an important role in integrating afferent synaptic activity with postsynaptic activity (Yuste and Denk, 1995). The geometry of a spine, in particular, the length of the spine neck and the size of the spine head are considered structural correlates of synapse function: While the length of the spine neck has been linked to the biochemical and electrical isolation of the spine compartment (Yuste et al., 2000; Yuste, 2013), the size of the spine head has been positively correlated with AMPA-receptor density, synaptic strength and spine stability (Matsuzaki et al., 2004; Kasai et al., 2010; McKinney, 2010). At the behavioral level, a critical role of spines in memory formation and cognition has been discussed (Segal, 2005; Bourne and Harris, 2007; Kasai et al., 2010; McKinney, 2010) and, indeed, a recent study has shown that spines are both necessary and sufficient for memory storage and memory trace formation (Abdou et al., 2018).

The function of spines does not only depend on the shape and size of their outer membranes but also on the molecular machinery within the spine subcompartment. A cellular organelle unique to spines is the spine apparatus (Gray, 1959; Spacek, 1985), which consists of the stacked endoplasmic reticulum (ER) and which modifies spine Ca^2+^ transients (Korkotian and Segal, 2011). Computational analyses showed that the geometry of the spine apparatus in relation to the geometry of the surrounding spine is a major determinant of second messenger dynamics (Cugno et al., 2018). Synaptopodin (SP) is an essential component of the spine apparatus and mice lacking SP do not form the organelle and show deficits in synaptic plasticity (Deller et al., 2003; Jedlicka et al., 2009; Vlachos et al., 2009; Vlachos et al., 2013; Zhang et al., 2013; Korkotian et al., 2014; Jedlicka and Deller, 2017) as well as in spatial learning (Deller et al., 2003).

The function of SP in spines is not limited to the formation of spine apparatus organelles. SP is also an actin-modulating protein, which affects actin microfilaments either directly by stabilizing F-actin (Mundel et al., 1997; Okubo-Suzuki et al., 2008) or indirectly via binding to alpha-actinin-2, Cdc42, RhoA or myosin V (Asanuma et al., 2005; Kremerskothen et al., 2005; Asanuma et al., 2006; Faul et al., 2007; Jedlicka and Deller, 2017; Konietzny et al., 2019). Since spine geometry primarily depends on actin assembly (Fischer et al., 1998; Matus, 2000; Konietzny et al., 2017), SP is likely to influence the geometry and long-term stability of spines. Indeed, short-term imaging experiments performed in dissociated cultures (Okubo-Suzuki et al., 2008; Vlachos et al., 2009) or acute slices (Zhang et al., 2013) implicated SP in plasticity-induced spine head expansion. We have now expanded on these previous studies and have used 2-photon time-lapse imaging of SP in granule cell spines to address the question of whether SP is important for spine head size and long-term spine stability.

## Materials and methods

### Animals

Adult male mice (12-34 weeks) lacking SP (SP-KO, C57BL/6J background; (Deller et al., 2003), wild-type mice (WT, C57BL/6J background) and Thy1-eGFP-SP x SP-KO mice (eGFP-SP-tg, C57BL/6J background (Vlachos et al., 2013) were used for the ex vivo analysis of granule cell spines in fixed slices. eGFP-SP-tg and SP-KO mice were used for the preparation of organotypic entorhino-hippocampal tissue cultures. eGFP-SP-tg mice were obtained by crossing homozygote Thy1-eGFP-SP transgenic mice (Vlachos et al., 2013) with SP-deficient mice (Deller et al., 2013). Thus, eGFP-SP-tg mice used for tissue culture preparation were monoallelic for eGFP-SP and devoid of endogenous SP. Adult mice used for fixed brain tissues were bred and housed at mfd Diagnostics GmbH, Wendelsheim, while mice used for organotypic entorhino-hippocampal tissue cultures were bred and housed at the animal facility of Goethe University Hospital Frankfurt. Animals were maintained on a 12 h light/dark cycle with food and water available ad libitum. Experimental procedures and animal care were performed in accordance with German animal welfare legislation and approved by the animal welfare officer of Goethe University Frankfurt, Faculty of Medicine. Every effort was made to minimize the distress and pain of animals.

### Intracellular injections of granule cells in fixed tissue

After delivery, animals were kept in an in-house scantainer for a minimum of 24 h. Animals were killed with an overdose of intraperitoneal Pentobarbital and subsequently intracardially perfused (0.1 M Phosphate Buffer Saline (PBS) containing 4% paraformaldehyde (PFA)). Tail biopsies were obtained after death to re-confirm the genotype. Brains were taken out immediately after perfusion, post-fixed (18 h, 4% PFA in 0.1 M PBS, 4° C), washed trice in ice-cold 0.1 M PBS, sectioned (250µm) on a vibratome (Leica VT 1000 S) and stored at 4 °C until use.

Intracellular injections of granule cells in fixed slices were performed as previously described (Germroth et al., 1989; Hick et al., 2015), with modifications. Hippocampal slices were placed in a custom-built, transparent and grounded recording chamber filled with ice-cold 0.1 M PBS. The chamber was attached to an epifluorescent microscope (Olympus BX51WI; 10x objective LMPlanFLN10x, NA 0.25, WD 21 mm) mounted on an x-y translation table (Science Products, VT-1 xy Microscope Translator). Sharp quartz-glass microelectrodes (Sutter Instruments, QF100-70-10, with filament) were pulled using a P-2000 laser puller (Sutter Instruments). Microelectrodes were tip-loaded with 0.75mM Alexa568-Hydrazide (Invitrogen) in HPLC-grade water (VWR Chemicals, HiPerSolv CHROMANORM) and subsequently back-filled with 0.1 M LiCl in HPLC-grade water. Microelectrodes were attached to an electrophoretic setup via a silver wire and 500 MΩ resistance. The tip of the microelectrode was navigated into the granule cell layer using a micromanipulator (Märzhäuser Wetzlar, Manipulator DC-3K). A square-wave voltage (1mV, 1 Hz) was applied using a voltage generator (Gwinstek SFG-2102). Granule cells were filled under visual control for at least 10 min or until no further labeling was observed. Injected sections were fixed (4% PFA in PBS, overnight, 4 °C, in darkness), washed in 0.1 M PBS and mounted on slides (Dako fluorescence mounting medium, Dako North America Inc.). Only granule cells with dendrites reaching the hippocampal fissure were used for analysis.

### Organotypic tissue cultures

Organotypic entorhino-hippocampal tissue cultures (300 µm thick) were prepared from postnatal 4-5 days old eGFP-SP-tg mice of either sex as previously described with minor modifications (Del Turco and Deller, 2007; Vlachos et al., 2012). Culture incubation medium contained 42% MEM, 25% Basal Medium Eagle, 25% heat-inactivated normal horse serum, 25 mM HEPES, 0.15% sodium bicarbonate, 0.65% glucose, 0.1 mg/ml streptomycin, 100 U/ml penicillin, 2 mM glutamax, adjusted to pH 7.30. The cultivation medium was refreshed every 2 to 3 days. All tissue cultures were allowed to mature in vitro for at least 25 days in a humidified incubator (5% CO2, at 35 °C).

### Laser microdissection

Tissue cultures used for laser microdissection were washed with 0.1 M PBS, embedded in tissue-freezing medium (Leica Microsystems) and shock-frozen (−80 °C) as previously described with minor modifications (Vlachos et al., 2013). Sections (10 μm) were cut on a cryostat (Leica CM 3050 S) and mounted on Poly Ethylene Terephthalate (PET) foil metal frames (Leicia Microsystems). Sections were fixed in acetone (−20 °C, 30 sec), stained with 0.1% toluidine blue (Merck) at room temperature for 20 sec before rinsing in ultrapure water (DNase/RNase free, Invitrogen) and dehydrated in 70% (vol/vol) and 100% ethanol. PET foil metal frames were mounted on a Leica LMD 6000 B system (Leica Microsystems) and a pulsed UV laser beam was directed along the borders of the dentate granule cell layer. Microdissected portions of the granule cell layer were collected in microcentrifuge tube caps placed underneath the frame. Caps contained guanidine isothiocyanate-containing buffer (RNeasy Mini Kit, Qiagen) and 1% β-mercaptoethanol (AppliChem GmbH). Microdissected tissue samples were stored at −80 °C until further processing.

### RNA extraction and quantitative PCR

Total RNA was isolated using the RNeasy Micro Plus Kit (Qiagen) and the purified RNA was reverse transcribed into cDNA with a High Capacity cDNA Reverse Transcription Kit (Applied Biosystems) following the manufacturers’ instructions as previously described with minor modifications (Vlachos et al., 2013). The cDNA was amplified using the StepOnePlus Real-Time PCR System (Applied Biosystems) with a standard amplification protocol (14 cycles of 95 °C for 15 sec and 60 °C for 4 min), and TaqMan PreAmp Master Mix Kit (Applied Biosystems) which contained 5 μL of PreAmp Master Mix, 2.5 μL of cDNA and 2.5 μL of Assay Mix (TaqMan Gene Expression Assays from Applied Biosystems; GAPDH, assay 4352932E; Synaptopodin, assay by design: forward primer, GTCTCCTCGAGCCAAGCA; reverse primer, CACACCTGGGCCTCGAT; probe, TCTCCACCCGGAATGC). Amplified cDNAs were diluted in ultrapure water (1:20) and quantitative PCR was performed using a standard amplification protocol (1 cycle of 50 °C for 2 min, 1 cycle of 95 °C for 10 min, 40 cycles of 95 °C for 15 sec and 60 °C for 60 sec; ending at 36 cycles).

### Adeno-associated virus (AAV) production

HEK293T cells were transfected with pDP1rs (Plasmid Factory), pDG (Plasmid Factory), and tdTomato-vector plasmid (12:8:5) by calcium phosphate seeding and precipitation (Grimm et al., 1998). Cells were collected 48 h after transfection, washed twice with 0.1 M PBS, centrifuged at 1.500 x g for 5 min and resuspended in 0.1 M PBS. Viral particles within the cells were released by 4 freeze-thaw cycles and the supernatant centrifuged at 3,200 x g for 10 min to remove cell debris. The final supernatant was collected, aliquoted and stored at −80 °C.

### Viral labelling

To label dentate granule cells, tissue cultures were transduced with an AAV serotype 2 (AAV2) containing the gene for tdTomato under the human Synapsin 1 promoter (Radic et al., 2017). Local injections were performed on cultures using an injection pipette pulled from thin-walled borosilicate capillaries (Harvard Apparatus, 30-0066). The pipettes were held by a head stage with a HL-U holder (Axon Instruments) and positioned using a micro-manipulator (Luigs & Neumann). Approximately 0.05-0.1 µl of AAV2-hSyn-tdTomato was injected directly into the suprapyramidal blade of the dentate gyrus using a syringe. Tissue cultures were visualized with an upright microscope (Nikon FN1) equipped with a camera and software (TrueChrome Metrics) using a 10x water immersion objective lens (Nikon Plan Fluor, NA 0.30), which allowed precise injection of the AAV2 into the DG. Injections were performed 2-3 days after the tissue cultures were prepared.

### Electron microscopy

Tissue cultures were fixed for 2 h in 0.1 M sodium cacodylate buffer (CB) containing 4% PFA, 4% sucrose, 15% picric acid and 0.5% glutaraldehyde. After four washes in CB, the fixed cultures were resliced to 40 µm. Following a wash with 0.1 M tris-buffered saline (TBS), free floating sections were treated with 0.1% NaBH4 (Sigma-Aldrich) in TBS for 10 min to remove unbound aldehydes. Sections were then blocked with 5% Bovine Serum Albumin (BSA) in TBS for 1 h at room temperature.

For detection of SP, sections were first incubated with rabbit anti-SP primary antibody (1:1000; Synaptic Systems) in 2% BSA and 0.1 M TBS for 18 h at room temperature followed by incubation with biotinylated goat anti-rabbit Immunoglobin G secondary antibody (1:200; Vector Laboratories, Burlingame, CA) in 2% BSA and 0.1 M TBS for 75 min at room temperature. After three washes with TBS, sections were incubated in avidin-biotin-peroxidase complex (ABC-Elite, Vector Laboratories) for 90 min at room temperature and treated with diaminobenzidine solution (Vector Laboratories) for 2-15 min at room temperature. Sections were then incubated for silver-intensification in 3% hexamethylenetetramine (Sigma-Aldrich), 5% silvernitrate (AppliChem) and 2.5% disodiumtetraborate (Sigma-Aldrich) for 10 min at 60 °C, followed by 0.05% tetrachlorogold solution (AppliChem) for 3 min and 2.5% sodium thiosulfate (Sigma-Aldrich) for 3 min with three washes after each step using distilled water.

After immunostaining, sections were washed in 0.1 M CB, treated with 0.5% OsO4 (Plano) in 0.1 M CB for 30 min, dehydrated in increasing concentrations of ethanol and then 1% uranyl acetate (Serva) in 70% ethanol for 60 min, before being embedded in Durcupan (Sigma Aldrich) for ultrathin sectioning (60 nm thickness). Sections were collected on single slot Formvar-coated copper grids and investigated using an electron microscope (Zeiss EM 900).

### Confocal microscopy of fixed hippocampal slices

Confocal imaging of fixed dendritic segments from identified, Alexa568-labeled dentate granule cells in the outer molecular layer (OML) of the suprapyramidal blade was done with an Olympus FV1000 microscope and a 60x oil-immersion objective (UPlanSApo, NA 1.35, Olympus) using FV10-ASW software with 5x scan zoom at a resolution of 1024 x 1024 pixels. To ensure localization in the OML, 3D image stacks of dendritic segments (0.25 µm z-axis step size) were taken at a distance of 10-50 µm from the hippocampal fissure. Crossing dendritic segments or branch points were avoided to facilitate spine attribution to a given segment.

### 2-Photon time-lapse imaging of tissue cultures

Live imaging of tdTomato-labelled granule cells with eGFP-tagged SP clusters was done using an upright 2-photon microscope (Scientifica MPSLSC-1000P, East Sussex, UK) equipped with a 40x water immersion objective (Zeiss Plan-Apochromat, NA 1.0) and a Ti-sapphire mode-locked laser (MaiTai, Spectra-Physics) tuned to a wavelength of 1000 nm to excite both eGFP and tdTomato. The membrane insert containing the cultures was placed in a petri dish containing warm imaging buffer consisting of 129 mM NaCl, 4 mM KCl, 1 mM MgCl2, 2 mM CaCl2, 4.2 mM glucose, 10 mM HEPES buffer solution, 0.1 mM Trolox, 0.1 mg/ml streptomycin, 100 U/ml penicillin, and pH 7.4. 3D image stacks of dendritic segments (∼50-80 µm in length) located in the middle to outer molecular layers (15-25 images per stack, 0.5 µm z-axis step size) were acquired using ScanImage 5.1 (Pologruto et al., 2003) with 8x digital zoom at a resolution of 512 x 512 pixels, i.e. 0.07 µm x 0.07 µm in the focal plane. The same dendritic segments across imaging sessions were identified using nearby landmarks and neighboring dendrites. Imaging time was minimized (only 1 dendritic segment per culture was imaged) to minimize the risk of phototoxic damage.

### Image processing and data analysis

The images obtained were deconvolved with Huygens Professional Version 17.10 (Scientific Volume Imaging, The Netherlands, http://svi.nl). Image processing and data analysis were then performed using Fiji version 1.52h (Schindelin et al., 2012), with spine analysis adapted from published criteria (Holtmaat et al., 2009).

Dendritic spines of all shapes were assessed manually on z-stacks of dendritic segments in the middle to outer molecular layers. Only protrusions emanating laterally in the x-y directions, not above or below the dendrite, and exceeding the dendrite for at least 5 pixels (0.35 µm; 0.2 µm for confocal) were included for analysis (Holtmaat et al., 2009; Vlachos et al., 2012).

For spine head size and SP cluster size measurements, the maximum cross-sectional area of the spine head or SP cluster in one of the x-y planes within the z-stack was measured. A spine was considered SP+ if the SP cluster overlapped with the spine head and/or neck in both the x-z and y-z directions when scrolling through the z-stacks. For spine stability analysis, we examined ∼15 µm long subsegments exhibiting minimal dendritic distortion over the total observation period. Individual spines were re-identified at consecutive points in time based on their relative positions to nearby landmarks and neighboring spines (Holtmaat et al., 2009; Vlachos et al., 2012). The presence or absence of a particular spine at a certain time point was verified by scrolling through the z-stacks to avoid misclassification due to rotation of the dendritic segment.

Spine turnover ratio representing the fraction of all observed spines that appear and/or disappear during the total imaging period was measured (adapted from Holtmaat, et al., 2009). Spine turnover ratio = (Ngained + Nlost) / (2 x Ntotal), where Ngained is the number of spines gained between the initial observation time point of t = −1 and the final observation time point of t = 12, Nlost is the number of spines lost between t = −1 and t = 12, and Ntotal is the total number of spines that were observed at one or more points in time. Spine formation ratio = (Ntotal – Ninitial) / (Ntotal), where Ninitial is the number of spines observed at t = −1. Spine loss ratio = (Ntotal – Nfinal) / (Ntotal), where Nfinal is the number of spines observed at t = 12.

### Statistical analysis

Statistical tests and n-values are indicated in figure captions. Statistical tests were chosen based on the experimental conditions, i.e. (1) matched-pairs, (2) two independent groups, (3) multiple matched-pairs, or (4) multiple independent groups. Thus, statistical comparisons were performed using Wilcoxon signed rank test with Pratts modification accounting for ties and zero differences in case of comparing integer values (differences between matched pairs under the null hypothesis of no difference), Mann-Whitney U-test (comparing two independent groups), Friedman test (for differences between matched pairs of multiple groups), and Kruskal-Wallis test (comparing multiple groups). Dunn’s multiple comparison test was applied following one of the two latter tests in case of significant P-values. All statistical tests were performed using GraphPad Prism 6. If *P* values were less than 0.05 the null-hypothesis was rejected. Statistical values were expressed as mean ± standard error of mean (SEM), unless otherwise stated. \**p* < 0.05, \*\**p* < 0.01, \*\*\**p* < 0.001.

### Modeling of spine survival and loss

The fractions of surviving spines observed at distinct time points were fitted to different models of spine survival and loss, respectively (see results). Curve fitting was performed with the user-defined fitting function of GraphPad Prism 6. Decay functions were fitted to the data weighted by 1/Y^2^ using least squares fit. The resulting spine loss curves correspond to the probability that single spines experience a particular survival time. Therefore, the loss curves represent the cumulative density function of spine survival times. We derived the distribution of spine survival times underlying the spine loss curves from the inverse loss curves. Since there is no analytical solution for the inverse of the conditional two stage decay, the inverse loss functions (i.e. the inverse cumulative density functions of spine survival times) were obtained from the spine loss models numerically as follows: According to the diverse loss models, the fractional survival was calculated for 10^7^ equally spaced time points ranging from zero to 30 times the longest time constant of the respective decay function. For each of 10^5^ equally spaced bins of fractional survival, the mean of the corresponding times (X-axis) was defined as the survival time corresponding to 1 minus the center of the respective fractional survival interval. The distribution of survival times displayed in the figures was derived as the histogram of these values using a bin size of 100. The median as well as the 10th, 25th, 75th, and 90th percentile of these distributions are also displayed in the figures. Statistical comparisons of median survival times between groups were based on the medians of 10 random samples of sizes equal to the number of observed spines per group using the Mann-Whitney U-Test. Derivation of survival times was performed using LabVIEW 2019 (National Instruments).

## Results

### Synaptopodin is present in large granule cell spines in vivo

Previous work reported a positive correlation between the plasticity-related protein SP and spine head size (Okubo-Suzuki et al., 2003; Vlachos et al., 2009; Zhang et al., 2013). Since SP-deficient mice show deficits in synaptic plasticity (Deller et al., 2003; Jedlicka 2009; Zhang et al., 2013; Grigoryan and Segal, 2016), we speculated that SP-deficient neurons might have smaller spine head sizes, providing a structural explanation for the plasticity phenotype. Accordingly, we analyzed and compared spine head size in hippocampal tissue sections of wildtype and SP-deficient mice (Deller et al., 2003) bred on the C57BL/6J genetic background. Surprisingly, Alexa 586-filled wildtype and mutant granule cells (Fig. 1A,B) showed similar average spine head sizes (Fig. 1C) and spine head size distributions (Fig. 1D). Next, we re-visited the relationship between SP and spine head size that had been suggested earlier (Okubo-Suzuki et al., 2003; Vlachos et al., 2009; Zhang et al., 2013). Using Alexa 568-injected granule cells from eGFP-SP-tg mouse brain (Fig. 1E; Vlachos et al., 2013), we first studied the fraction of SP-positive spines and found SP clusters in ∼14% of spines (Fig. 1F). Spines containing SP (SP+) were significantly larger than spines without SP (SP-) (Fig. 1G) and showed a right shifted cumulative frequency distribution (Fig. 1H). In addition, SP cluster size was positively correlated with spine head size (Fig. 1I), demonstrating that large SP clusters are typically found in large spines. We conclude from these in vivo observations that SP is indeed tightly correlated with spine size. However, SP does not appear to be a major regulator of spine head size, since neither average spine head size nor spine head size distributions were affected by SP-deficiency (Fig. 1D).

**Figure 1.**
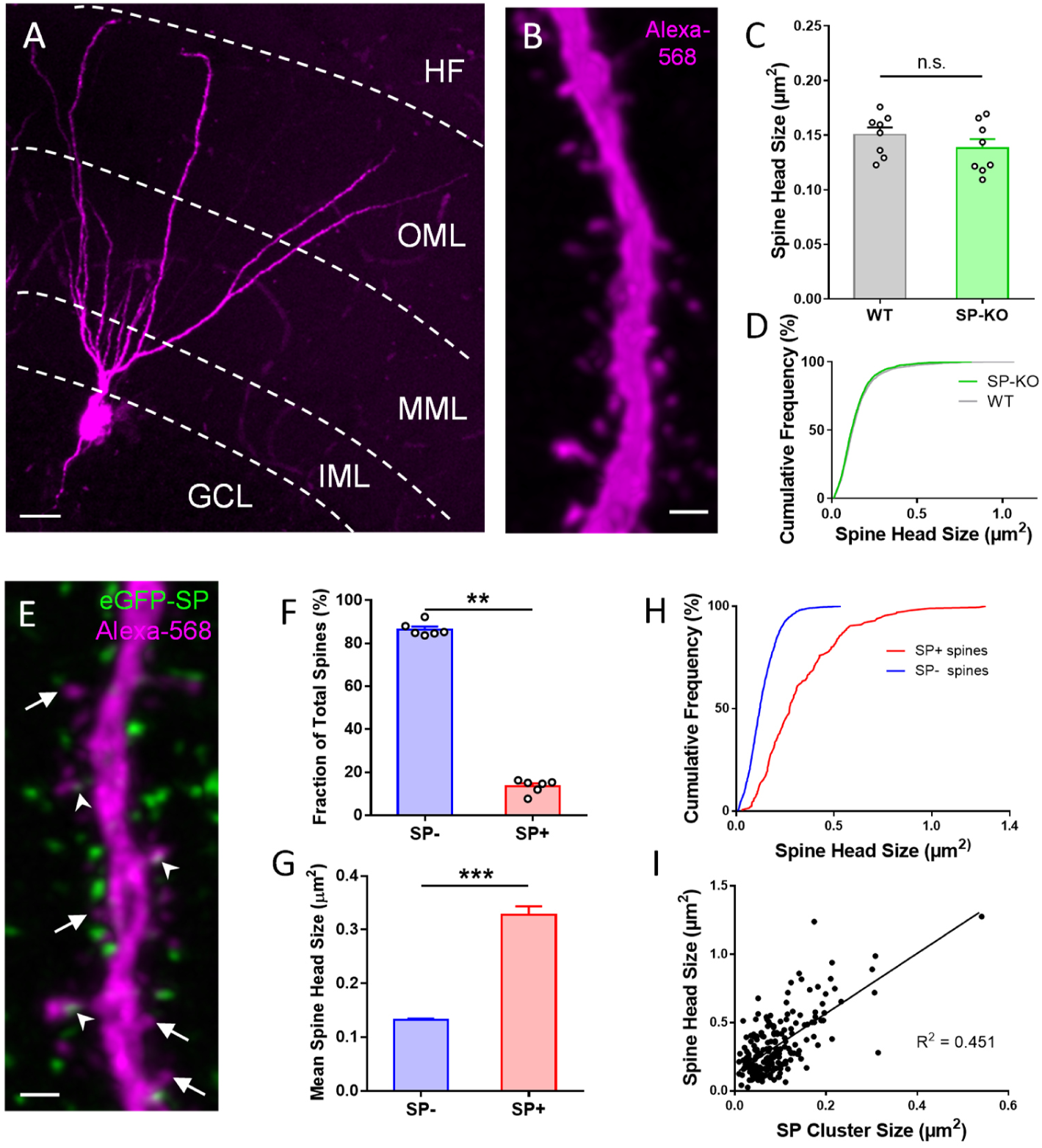
SP is associated with large granule cell spines in adult mouse dentate gyrus. (A) Granule cell located in the suprapyramidal blade of the dentate gyrus of a SP-knock-out (SP-KO) mouse intracellularly filled with the fluorescent dye Alexa-568 (magenta; fixed tissue). Dendritic segments in the outer molecular layer (OML) were used for analysis. MML: middle molecular layer. IML: inner molecular layer. GCL: granule cell layer. HF: hippocampal fissure. Scale bar = 20 µm. (B) Dendritic segment of a SP-KO granule cell shown at higher magnification. Scale bar = 1 µm. (C) Spine head sizes of wildtype (WT) and SP-KO mice. n.s., not significant; p = 0.291, Mann-Whitney U-test. WT mice and SP-KO mice, n = 8 per group (3 dendritic segments per animal; 1885 WT spines, 2158 SP-KO spines). (D) Cumulative frequency plots of spine head sizes of wildtype (grey) and SP-KO (green) mice. (E) Granule cell located in the suprapyramidal blade of the dentate gyrus of a Thy1-eGFP-SP-transgenic mouse bred on a SP-KO background (eGFP-SP-tg mouse) intracellularly filled with Alexa-568 (magenta; fixed tissue; OML). Arrowheads point to eGFP-SP clusters (green) in SP-positive (SP+) dendritic spines. Arrows mark SP-negative (SP-) spines. Scale bar = 1 µm. (F) Fractions of SP- (∼86.5%) and SP+ (∼13.5%) spines. **p < 0.0022, Mann-Whitney U-test. eGFP-SP-tg mice, n = 6. (G) Mean spine head size of SP- (∼0.133 µm^2^) and SP+ (∼0.328 µm^2^) spines. ***p < 0.0001, Mann-Whitney U-test. SP+ spines n = 200; SP- spines n = 1497. (H) Cumulative frequency plots of spine head sizes of SP- (blue) and SP+ (red) spines. SP+ spines = 200; SP- spines = 1497. (I) Correlation analysis of spine head size and SP cluster size: Spearman coefficient of correlation = 0.536, 95% C.I.: 0.426 – 0.630. ***p < 0.0001. Linear regression analysis: R^2^ = 0.451. n = 200 spines.

This surprising finding raised the question of what is the role of SP in large spines, if SP is not a regulator of head size? Since SP is an actin-modulating plasticity-related protein, it has been speculated that SP could also regulate spine stability (e.g. Deller et al., 2000).

As previous reports have shown that large cortical spines are more stable than small spines (Kasai et al., 2003; Matsuzaki et al., 2004; Bourne and Harris, 2008; Kasai et al., 2010; McKinney, 2010), we speculated that the presence of SP in these spines could explain or at least contribute to their higher stability. To test this hypothesis, we used an organotypic tissue culture preparation, which allowed us to follow not only spine geometry – which can also be done in vivo – but also the dynamics of SP clusters within spines over time.

### Synaptopodin is present in large granule cell spines in organotypic tissue cultures

Organotypic tissue cultures of the entorhinal cortex and hippocampus were generated from eGFP-SP- tg mice. In these cultures SP clusters were abundant in the molecular layer of the dentate gyrus (Fig. 2A), similar to what has been described for wildtype animals after SP immunolabeling (Deller et al., 2000b; Bas Orth et al., 2005). To visualize single granule cells and their spines, tissue cultures were virally transduced on 2-3 DIV using an AAV2-tdTomato virus. After approximately three weeks in vitro, tdTomato expressing granule cells were imaged and spines with SP (SP+) and without SP (SP-; Fig. 2B) were identified. SP+ spines also contained SP-positive spine apparatus organelles (Fig. 2C). SP mRNA expression levels in eGFP-SP-tg cultures were similar to SP mRNA expression levels in granule cells of SP +/+ wildtype cultures (Fig. 2D). Since SP had been suggested to regulate spine head size (see previous paragraph), we also investigated average spine head size and spine head size distribution in SP-deficient cultures (Fig. 2E, F). No significant differences between SP-deficient and eGFP-SP-tg granule cells were seen, in line with our in vivo observations (c.f. Fig. 1), suggesting again that SP is not a major regulator of spine head size.

**Figure 2.**
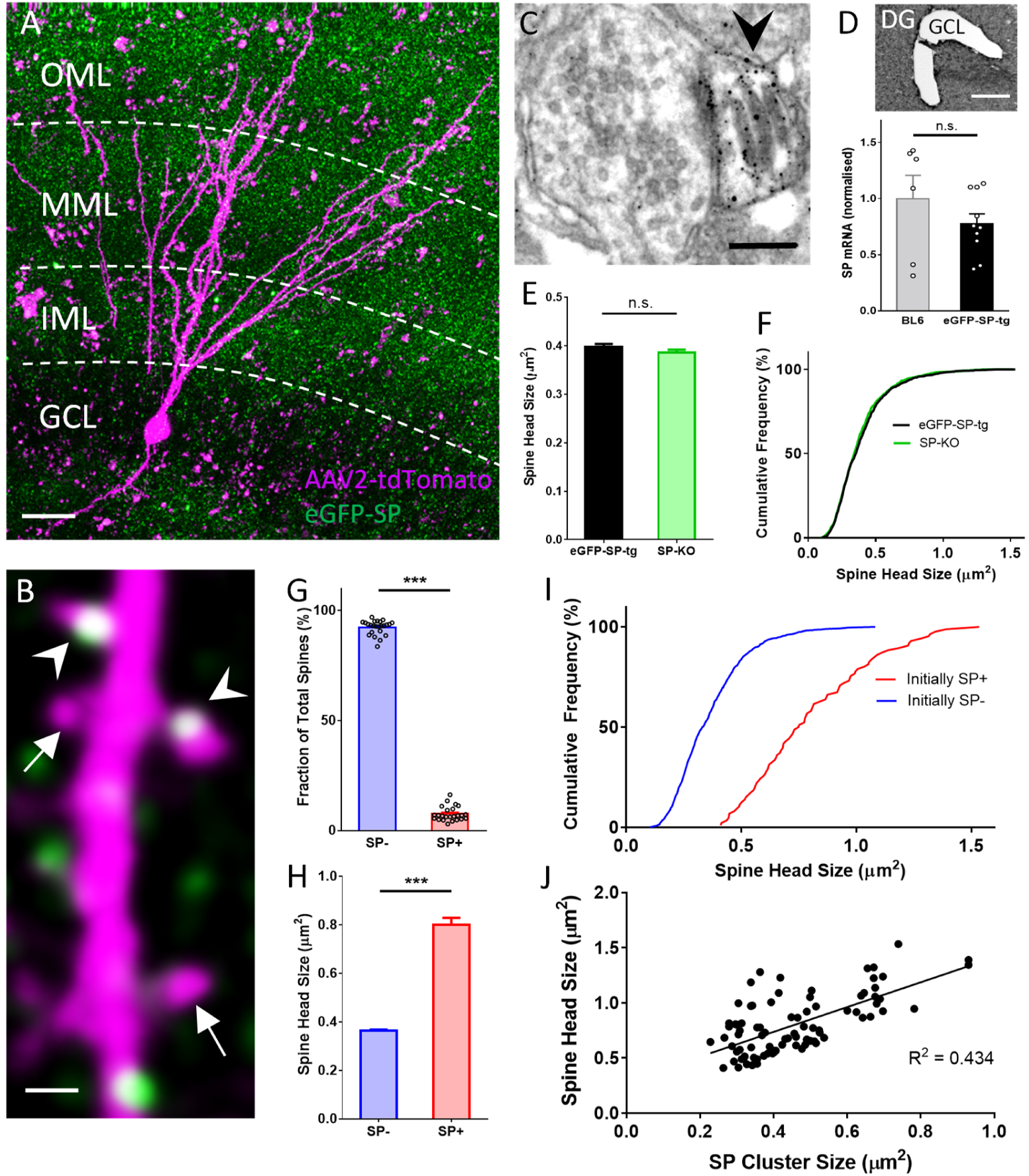
SP in dendritic spines of granule cells in organotypic tissue cultures of hippocampus. (A) Granule cells in organotypic entorhino-hippocampal tissue cultures (OTCs) of eGFP-SP-tg (green) mice were virally transduced (AAV2) with tdTomato (magenta) at day in vitro (DIV) 2-3. Dendritic segments located in the middle molecular layer (MML) or in the outer molecular layer (OML) were imaged. GCL, granule cell layer; IML, inner molecular layer. Scale bar = 20 µm. (B) Single plane 2-photon image of a granule cell dendrite in the OML. Arrowheads point to SP+ spines; arrows indicate SP-spines. Scale bar = 1 µm. (C) Electron micrograph of a SP+ spine (arrowhead) containing an immunolabeled spine apparatus in an OTC from an eGFP-SP-tg mouse. Scale bar = 0.2 µm. (D) SP-mRNA levels in microdissected granule cell layers (GCL; upper panel) from OTCs of eGFP-SP-tg and C57BL/6J (BL6) wildtype mice were not significantly different (lower panel). p = 0.353, Mann-Whitney U-test. Number of BL6 tissue cultures n = 6; number of eGFP-SP-tg tissue cultures n = 10. DG, dentate gyrus. Scale bar = 100 μm. (E) Mean spine head sizes of eGFP-SP-tg (black) and SP-KO (green) granule cells. n.s., not significant. p = 0.211, Mann-Whitney U-test. Number of spines: eGFP-SP-tg n = 1107; SP-KO n = 1113, obtained from 24 segments, 1 segment per culture. (F) Cumulative frequency plots showing the distribution of spine head sizes of these spines. (G) SP+ spines comprised 7.6% of the total spine population. ***p < 0.0001, Mann-Whitney U-test. Percentage per eGFP-SP-tg dendritic segment; n = 24 segments. (H) SP+ spines were significantly larger than SP-spines. ***p < 0.0001, Mann-Whitney U-test. Number of SP+ spines n = 87, SP- spines n = 1021. (I) Cumulative frequency plots showing the distribution of spine head sizes of SP+ and SP- spines. (J) SP cluster size and spine head size of SP+ spines are tightly correlated: Spearman coefficient of correlation = 0.565, 95% C.I.: 0.396 – 0.697. ***p < 0.0001. Linear regression analysis: R^2^ = 0.434. n = 87 spines. The following figure supplements are available for Figure 2. **Figure S1 related to Figure 2. Measurement of dendritic spine head size and SP cluster size.**

Next, we analyzed the fraction of SP+ and SP- spines and found that approximately 8% of all granule cell spines are SP+ (Fig. 2G). Analysis of head sizes of SP+ and SP- spines (Fig. 2; Suppl. Fig 1 related to Fig. 2) revealed that SP+ spines are on average much larger than SP- spines (SP- spines: 0.364 ± 0.005 µm^2^; SP+ spines: 0.801 ± 0.029 µm^2^; Fig. 2H), similar to what was observed in vivo (c.f. Fig. 1).

Likewise, cumulative frequency diagrams showed that the population of SP+ spines is right-shifted towards larger spine head sizes compared to SP- spines (Fig. 2I). Finally, correlation analysis of SP cluster size and spine head size demonstrated a positive correlation between the two parameters (Fig. 2J). We conclude from these observations that with regard to SP granule cells in organotypic tissue cultures are highly similar to granule cells in vivo (c.f. Fig. 1). Furthermore, we conclude that SP is preferentially found in large granule cell spines.

### Presence and size of Synaptopodin clusters in vitro are tightly associated with bidirectional changes in spine head size

After demonstrating significant differences in average spine head size between SP+ and SP- spines, we wondered whether this relationship between SP and spine head sizes is also seen under dynamic conditions, i.e. when spines undergo changes in their head size. For this, we studied how insertion and loss of SP clusters from spines affects spine head geometry (Fig. 3). Single spines containing SP and spines devoid of SP were identified and spine head sizes were measured on two consecutive days (Fig. 3). Four groups of spines were distinguished: Spines gaining SP (Fig. 3A), spines losing SP (Fig. 3B), spines maintaining SP (Fig. 3C), and spines remaining SP- (Fig. 3D). This analysis revealed a robust correlation between increases in spine head size and insertion of SP and decreases of spine head size and loss of SP from spines (Fig. 3).

**Figure 3.**
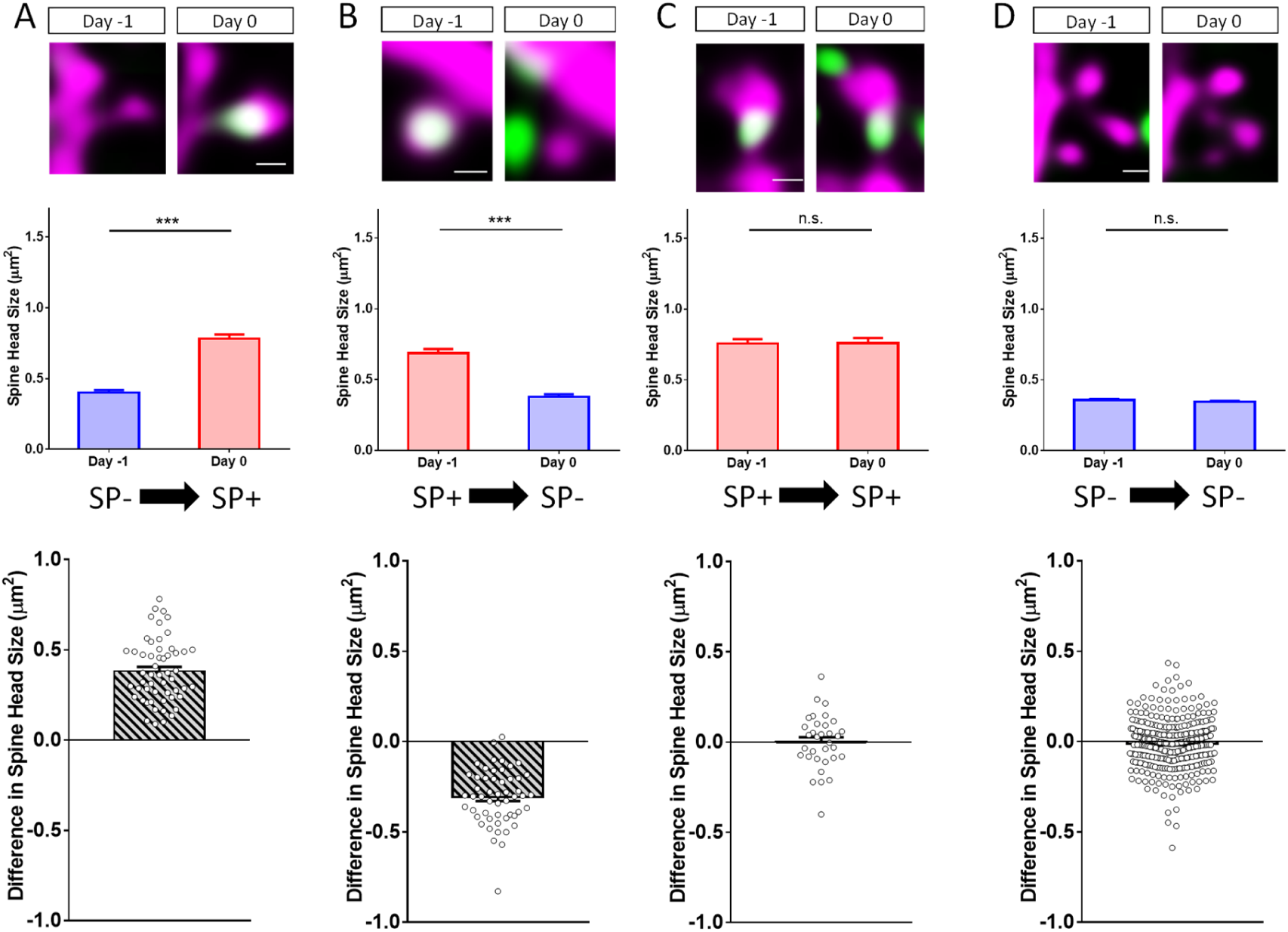
Changes in spine SP content are associated with bidirectional changes in spine head size. Sample spines (upper panels), mean spine head size changes (middle panels) and differences in spine head size of individual spines (lower panels) are illustrated. (A) Example of a spine that became SP+ between day −1 and 0. The mean maximum cross-sectional spine head area of spines of this group increased significantly. ***p < 0.0001, Wilcoxon matched-pairs signed rank test. Number of spines n = 54. Scale bar = 0.5 µm. (B) Example of a spine that was SP+ and became SP- between day −1 and 0. The mean maximum cross-sectional spine head area of spines of this group decreased significantly. ***p < 0.0001, Wilcoxon matched-pairs signed rank test. Number of spines n = 52. Scale bar = 0.5 µm. (C) Example of a spine that remained SP+. The mean cross-sectional spine head area of spines of this group did not change significantly. p = 0.850, Wilcoxon matched-pairs signed rank test. Number of spines n = 33. Scale bar = 0.5 µm. (D) Example of a spine that remained SP-. The mean cross-sectional spine head area of spines of this group did not change significantly. p = 0.060, Wilcoxon matched-pairs signed rank test. Number of spines n = 340. Scale bar = 0.5 µm.

### The presence of Synaptopodin in spines is associated with long-term spine survival

The stability of spines is important for maintaining specific connections between neurons. Since SP stabilizes F-actin (Okubo-Suzuki et al., 2008), we speculated that SP could have an effect on long-term spine stability (Deller et al., 2000a; Jedlicka and Deller, 2017).

To address this hypothesis, we used time-lapse imaging and followed single spines for two weeks (Fig. 4A). During this time period, we determined their SP content, head sizes and fate (Fig. 4A, B). This analysis revealed major differences in the survival time of SP+ and SP- spines (Fig. 4C-E): Whereas the median survival time of SP+ spines was ∼17.5 days, the median survival time of SP- spines was only ∼6.8 days (Fig. 4C).

**Figure 4.**
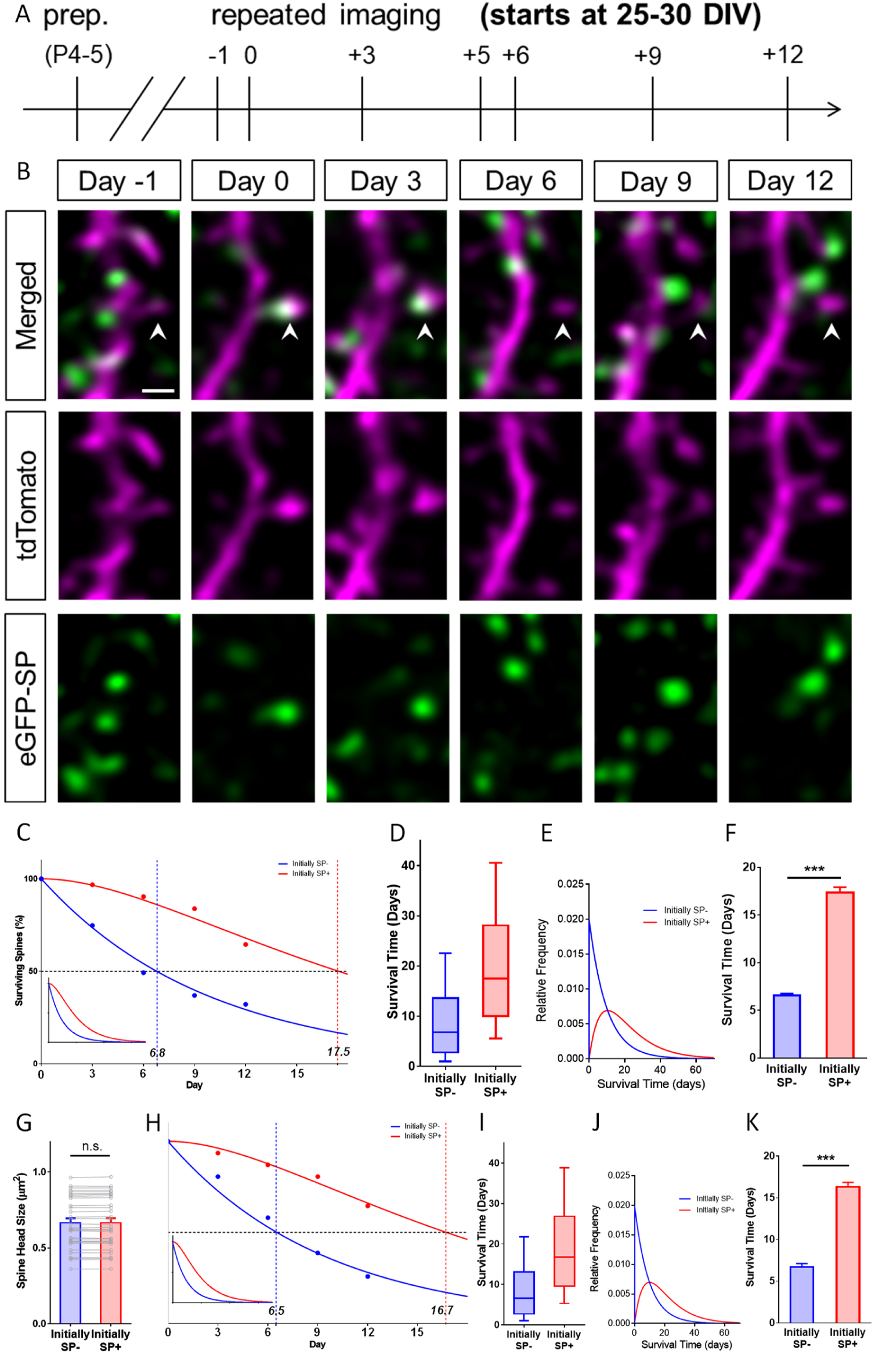
SP+ spines are highly stable spines. (A) Schematic of the experimental design. OTCs were prepared at postnatal day (p) 4-5 from eGFP-SP-tg mice and allowed to mature for 25-30 DIV. Time-lapse 2-photon imaging of identified dendritic segments was performed at time points as indicated. (B) Time-lapse imaging of a granule cell dendrite over 14 days illustrates the dynamics of SP clusters within individual spines. Arrowhead points to a spine that became SP+ on day 0, stayed positive on day 3, lost SP between days 3 and 6 and then remained SP- until day 12. Scale bar = 1 µm. (C) The survival of SP+ (red curve) and SP- spines (blue curve) was studied from day 0 until day 12. Dots: observed fractions; curves: fitted to observation points. The median survival of initially SP- spines was ≈ 6.8 days whereas the median survival of initially SP+ spines was ≈ 17.5 days. Number of spines at day 0: SP+ = 31, SP- = 392. Inset: Calculated decay curves until day 70. SP- spines followed a single stage decay function whereas SP+ spines followed a conditional two stage decay function. (D) Survival times of the total populations of SP+ and SP- spines derived from the fitted survival curves. Boxes indicate median, lower and upper quartiles. Whiskers indicate 10th and 90th percentiles, respectively. (E) Relative frequency distributions of these survival times. (F) The survival time of SP+ spines was significantly longer than SP- spines; computational model; ***p < 0.0001, Mann-Whitney U-test; based on 10 samples of 31 SP+ and 392 SP- spines. (G) To control for differences in spine head size, SP+ and SP- spines of equal head size were matched. The spine head size of these pairs was not significantly different. p = 0.310, Wilcoxon matched-pairs signed rank test. Number of size-matched pairs n = 31. (H) The decay curve of these size-matched spines was very similar to the decay curve of all SP+ and SP- spines. The median survival of SP- spines of this population was ≈ 6.5 days, whereas size-matched SP+ spines showed a median survival of ≈ 16.7 days. Inset: Calculated decay curves until day 70. (I) Survival times of the total populations of SP+ and SP- spines derived from the fitted survival curves. (J) Relative frequency distributions of these survival times. (K) The survival time of size-matched SP+ spines was significantly higher than that of SP- spines; computational model; ***p < 0.0001, Mann-Whitney U-test. based on 10 samples of 31 SP+ and 31 SP- spines.

In addition to this notable difference in median survival time, the curves that fitted our data turned out to be fundamentally different: Whereas SP- spines followed a single phase exponential decay function, the decay of SP+ spines could only be fitted using a conditional double phase exponential decay function (see below; Fig. 4C). Based on these two functions the distributions of long-term survival times (Fig. 4D, E) of the SP- and SP+ spine populations were calculated (see methods). Finally, a data-driven simulation of spine survival was performed to test whether the survival times of the two spine populations are different. This approach revealed – within the parameters used – that SP- and SP+ spines differ significantly in their survival times (Fig. 4F).

Since SP+ spines are larger than SP- spines (c.f. Fig. 1G, 2H) the observed effect on spine survival could simply be the result of differences in head size between the two groups. As large spines have been shown to be more stable (Kasai et al., 2003; Matsuzaki et al., 2004; Bourne and Harris, 2008; Kasai et al., 2010; McKinney, 2010), this would be a straightforward explanation. We controlled for this possibility by comparing the survival times of SP+ and SP- spines of equal head size (matched pairs; Fig. 4G). Even under these highly constrained conditions the difference in spine survival between SP+ and SP- spines was still observed (Fig. 4H). In line with this, the overall distributions (Fig. 4I), the relative frequency distributions of spine survival times of the calculated total populations of spines (Fig. 4J), as well as simulations of spine survival (Fig. 4K) were comparable to those found for all spines, i.e. non-size-matched spines.

### Synaptopodin increases survival of small, medium and large spines

Since the stability of large spines is higher than the stability of small spines (Kasai et al., 2003; Matsuzaki et al., 2004; Bourne and Harris, 2008; Kasai et al., 2010; McKinney, 2010), we wondered whether this difference could be caused by the presence of SP within large spines. This appeared to be a possibility since SP is tightly associated with the population of large and stable spines and rare in the group of small and less stable spines (Fig. 5A, B). Should the increased stability of large spines depend on SP we made two testable predictions: (1) If SP is present in small spines, it should increase survival of these spines irrespective of their size, and, (2) if SP is absent from large spines, these spines should be pruned faster in spite of their large size. To test these hypotheses, we subdivided SP+ and SP- spines into four size classes (Fig 5B; very small, small, medium and large spines) and compared the survival times of SP- and SP+ spines belonging to each of these classes. To test the first hypothesis, we analyzed the three smaller size groups, which comprise the majority (∼95%) of all spines (Fig. 6A). SP was completely absent from the very small spine group, in which ∼41% of all SP- spines are found (Fig. 5B). This spine population rapidly decayed with time following a single exponential decay function (Fig. 5C-F).

**Figure 5.**
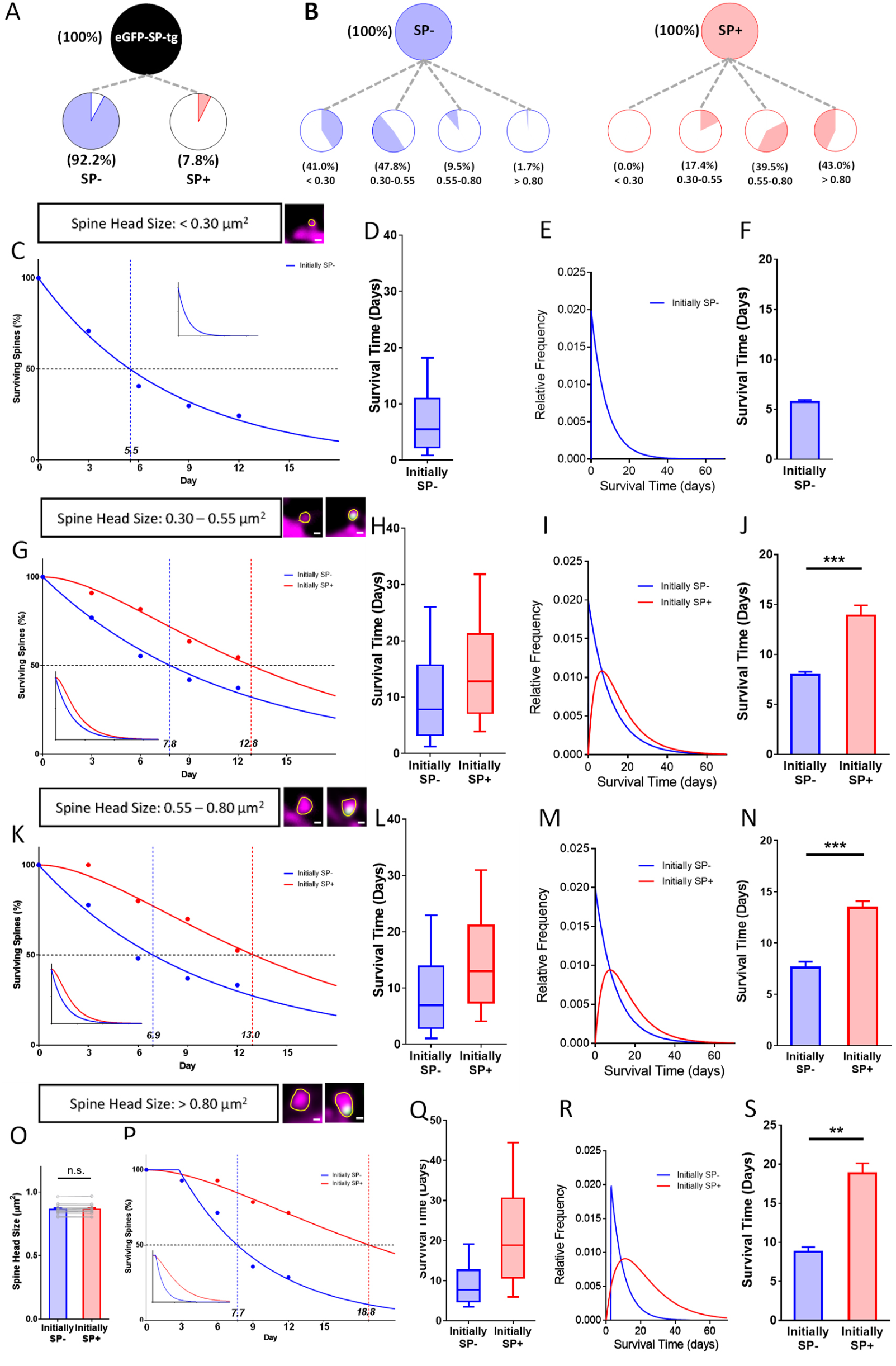
Presence of SP in spines determines their long-term survival. (A) Fraction of SP- (≈ 92.2%) and SP+ spines (≈ 7.8%) in eGFP-SP-tg granule cells on day 0. Number of SP+ spines = 87, SP- spines = 1021, obtained from entire imaged segments; number of cultures: eGFP-SP-tg = 24. (B) SP- (blue) and SP+ spines (red) were divided into four size classes: very small (< 0.3µm^2^), small (0.3-0.55 µm^2^), medium (0.55-0.8 µm^2^) and large (> 0.8 µm^2^) spines. Whereas most SP- spines were found in the very small (41.0%) and small (47.8%) spine groups, the majority of SP+ spines were in the medium (39.5%) and large (43.0%) size groups. (C) Survival curve of very small spines (example shown) from day 0 until day 12. Dots: observed fractions; curves: fitted to observation points. SP- spines (blue line) followed a single exponential decay curve (median survival ≈ 5.5 days). Inset: extrapolated decay curve until day 70. Number of very small SP- spines = 148. (D) Survival time of the total population of SP- spines derived from the fitted survival curve. Box indicates median, lower and upper quartiles. Whiskers indicate 10th and 90th percentiles, respectively. (E) Relative frequency distribution of very small SP- spines. (F) Mean survival time of very small SP- spines; computational model; 10 samples, 148 SP- spines. (G) Survival curves of small SP- and SP+ spines (examples shown). SP- spines (blue line; median survival ≈ 7.8 days) followed a single exponential decay curve, SP+ spines (red line; median survival ≈ 12.8 days) followed a conditional two-stage decay curve. Inset: extrapolated decay curves until day 70. Number of SP- spines = 217; SP+ spines = 11. (H) Survival times and (I) relative frequency distributions of small SP- and SP+ spines. (J) Mean survival time of SP+ spines compared to SP- spines; computational model. ***p = 0.0003, Mann-Whitney U-test. 10 samples; 11 SP+ and 217 SP- spines. (K) Survival curves of medium sized SP- and SP+ spines (examples shown). SP- spines (blue line; median survival ≈ 6.9 days) followed a single exponential decay curve, SP+ spines (red line; median survival ≈ 12.9 days) followed a conditional two-stage decay curve. Inset: extrapolated decay curves until day 70. Number of SP- spines = 27; SP+ spines = 40. (L) Survival times and (M) relative frequency distributions of medium SP- and SP+ spines. (N) Mean survival time of SP+ spines compared to SP- spines; computational model. ***p < 0.0001, Mann-Whitney U-test. 10 samples; 40 SP+ and 27 SP- spines. (O) SP+ and SP- spines were size-matched in the largest spine group (mean spine head sizes; not significant; p = 0.475, Wilcoxon matched-pairs signed rank test). Number of size-matched pairs n = 14. (P) Survival curves of size-matched large SP- and SP+ spines (examples shown). SP- spines (blue line; median survival ≈ 7.7 days) followed a single exponential decay curve, SP+ spines (red line; median survival ≈ 18.8 days) followed a conditional two-stage decay curve. Inset: calculated decay curves until day 70. (Q) Survival times and (R) relative frequency distributions of large SP- and SP+ spines. (S) Mean survival time of SP+ spines compared to SP- spines; computational model. ***p < 0.0001, Mann-Whitney U-test. 10 samples; 14 SP+ and 14 SP- spines.

**Figure 6.**
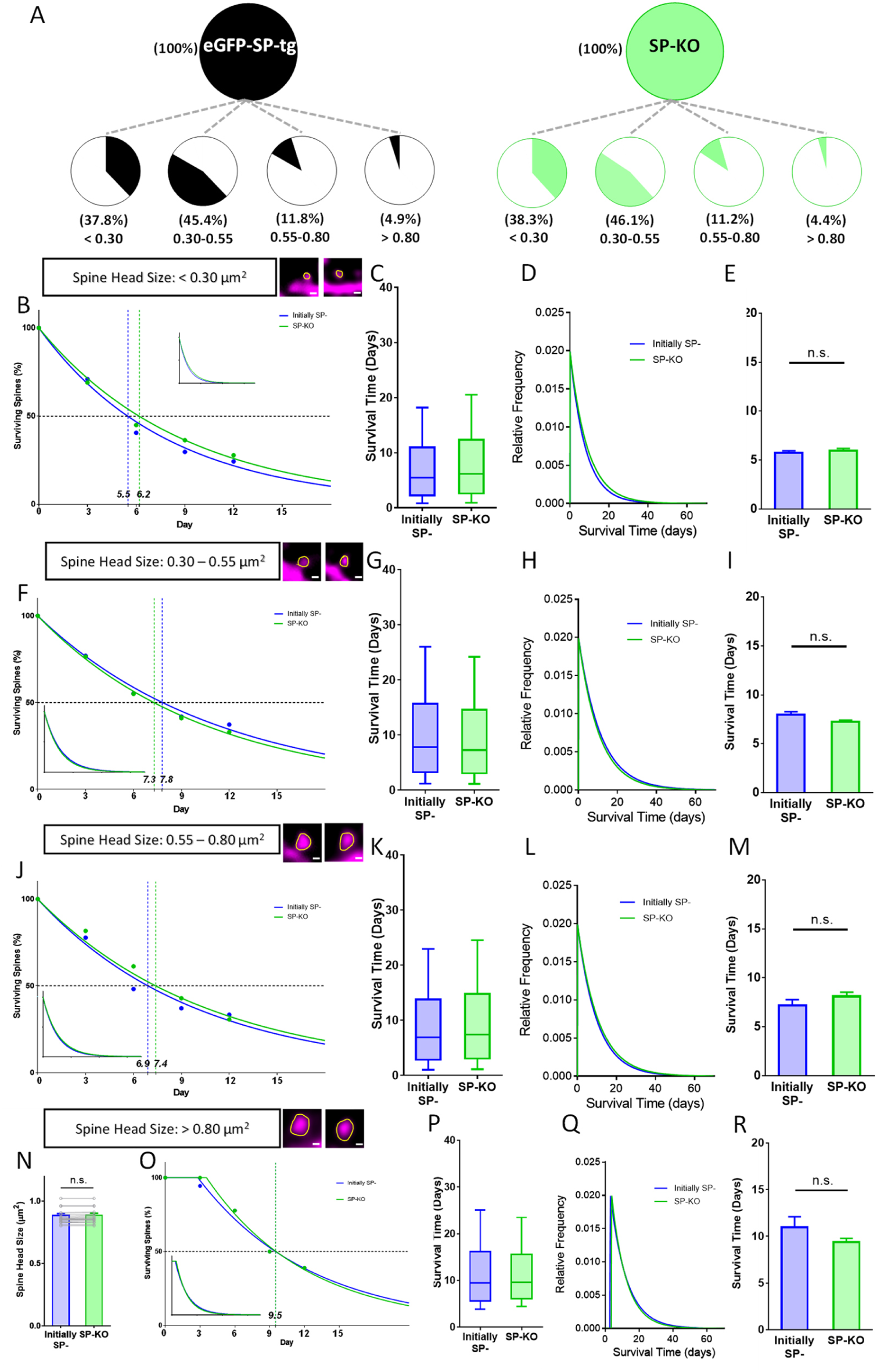
Spines of SP-deficient mice decay like SP- spines. (A) Spine head size distributions of granule cell dendrites of eGFP-SP-tg and SP-KO mice were similar. Number of spines: eGFP-SP-tg = 1107; SP-KO = 1113, obtained from entire imaged segments; number of cultures: eGFP-SP-tg = 24; SP-KO = 21. (B) Survival curve of very small spines (examples shown) from day 0 until day 12. Dots: observed fractions; curves: fitted to observation points. SP- spines (blue line; median survival ≈ 5.5 days) and spines of SP-KO-mice (green line; median survival ≈ 6.2 days) followed single exponential decay curves. Inset: calculated decay curves until day 70. Number of SP- spines = 148; SP-KO spines = 187. (C) Calculated survival times and (D) relative frequency distributions of very small SP- and SP-KO spines. (E) Mean survival time of very small SP- and SP-KO spines; computational model. p = 0.529, Mann-Whitney U-test. 10 samples; 148 SP-, 187 SP-KO spines. (F) Survival curves, (G) calculated survival times, and (H) relative frequency distributions of small SP- and SP-KO spines. SP- spines (blue; median survival ≈ 7.8 days) and SP-KO spines (green, median survival ≈ 7.4 days) followed single exponential decay curves. Number of SP- spines = 217; SP-KO spines = 200. (I) Mean survival times; computational model. p = 0.089, Mann-Whitney U-test. 10 samples; 217 SP-, 200 SP-KO spines. (J) Survival curves, (K) calculated survival times, and (L) relative frequency distributions of medium-sized SP- and SP-KO spines. SP- spines (blue; median survival ≈ 6.9 days) and SP-KO spines (green, median survival ≈ 7.4 days) followed single exponential decay curves. Number of SP- spines = 27; SP-KO spines = 49. (M) Mean survival times; computational model. p = 0.280, Mann-Whitney U-test. 10 samples; 27 SP-, 49 SP-KO spines. (N) SP+ and SP-spines were size-matched in the largest spine group (mean spine head sizes; not significant, p = 0.109, Wilcoxon matched-pairs signed rank test. Number of size-matched pairs n = 18. (O) Survival curves, (P) calculated survival times, and (Q) relative frequency distributions of large SP- and SP-KO spines. SP-spines (blue; median survival ≈ 9.5 days) and SP-KO spines (green, median survival ≈ 9.5 days) followed single exponential decay curves. Number of size-matched pairs = 18. (R) Mean survival times; computational model. p = 0.393, Mann-Whitney U-test. 10 samples; 18 SP- and 18 SP-KO spines. The following figure supplements are available for Figure 6. **Figure S2 related to Figure 6. SP-KO mice compensate for the loss of SP with an increased spine formation and an increased turnover ratio.**

In the small spine group ∼47.8% of all SP- and ∼17.4% of all SP+ spines are found (Fig. 5B), with SP+ spines representing ∼5% of all spines in this group. Compared to the SP- spines, which followed a single exponential decay curve, SP+ spines showed a longer survival time and followed a conditional double phase exponential decay function (Fig. 5G-J). In the group of medium-sized spines ∼9.5% of SP- and ∼39.5% of SP+ spines are found, with SP+ spines representing ∼60% of the spines in this size group. Again, SP- spines followed a single exponential decay curve, whereas SP+ spines showed longer survival times and their decay was compatible with a conditional double phase exponential decay function (Fig. 5K-N). We conclude from these data that small and medium-sized spines survive longer and follow a conditional double-phase decay kinetic if SP is present.

Finally, we looked at the population of large spines. Although these spines only account for ∼4.9% of all spines, they are considered especially important in the context of memory storage (Segal, 2005; Bourne and Harris, 2007; Kasai et al., 2010; McKinney, 2010; Abdou et al., 2018). In this group of spines only ∼1.7% of the SP- spines are found whereas ∼43% of SP+ spines are part of this subgroup. SP+ spines represent ∼65% of the spines in this size group. Comparing spine pairs of equal head sizes (Fig. 5O) major differences in their survival and decay kinetics were revealed: Whereas SP-spines followed a single exponential function, SP+ spines were best modelled using the conditional double-phase decay function (Fig. 5P-S). We conclude that large SP- spines are pruned fast. Their survival is comparable to that of very small (Fig. 5C), small (Fig. 5G) and medium sized (Fig. 5K) spines, suggesting that it is the presence of SP within spines rather than spine size per se which determines long-term spine survival.

### Synaptopodin-deficiency alters spine survival

We then turned to SP-deficient mice to study spine survival in the absence of SP. (Fig. 6). First, we compared the distribution of spines of the eGFP-SP-tg and the SP-KO mice and found comparable fractions of spines in the four size categories (Fig. 6A). This is in line with our observation that SP- deficiency does not affect average spine head size or spine head distribution in vivo (c.f. Fig. 1C, D) or in vitro (c.f. Fig. 2E, F). Next, we compared SP-deficient spines to SP- spines and found that these two groups of spines behave very similar with regard to their survival times and decay kinetics in each of the four size groups (Fig. 6B-R): survival times were similar and all curves followed single exponential decay functions. This demonstrates that the “spine phenotype” associated with SP-deficiency does not result in altered spine geometry but in altered spine survival, i.e. spine stability.

We then wondered how the lack of the stable SP+ spine population affects spine dynamics, in particular spine turnover, of the SP-deficient animals.

We reasoned that the reduced stability of large spines will have to be homeostatically compensated by an increased formation of spines and an increased spine turnover. Indeed, these dynamic parameters were significantly increased (Suppl. Fig. 2 related to Fig. 6), revealing a yet unknown phenotype of the SP-deficient mutants.

### Spines shrink and pass through a SP-negative stage before pruning

After investigating the survival of individual spines considering only their initial SP content (SP+ or SP-; previous paragraphs), we determined changes of the SP+ and SP- spine populations with regard to survival, head size and SP content (Fig. 7). For this, all spines were identified on day 0 as being either SP+ or SP-. For each of the following observation time points, these spines were categorized as either SP+, SP- or “lost” (Fig. 7A, B).

**Figure 7.**
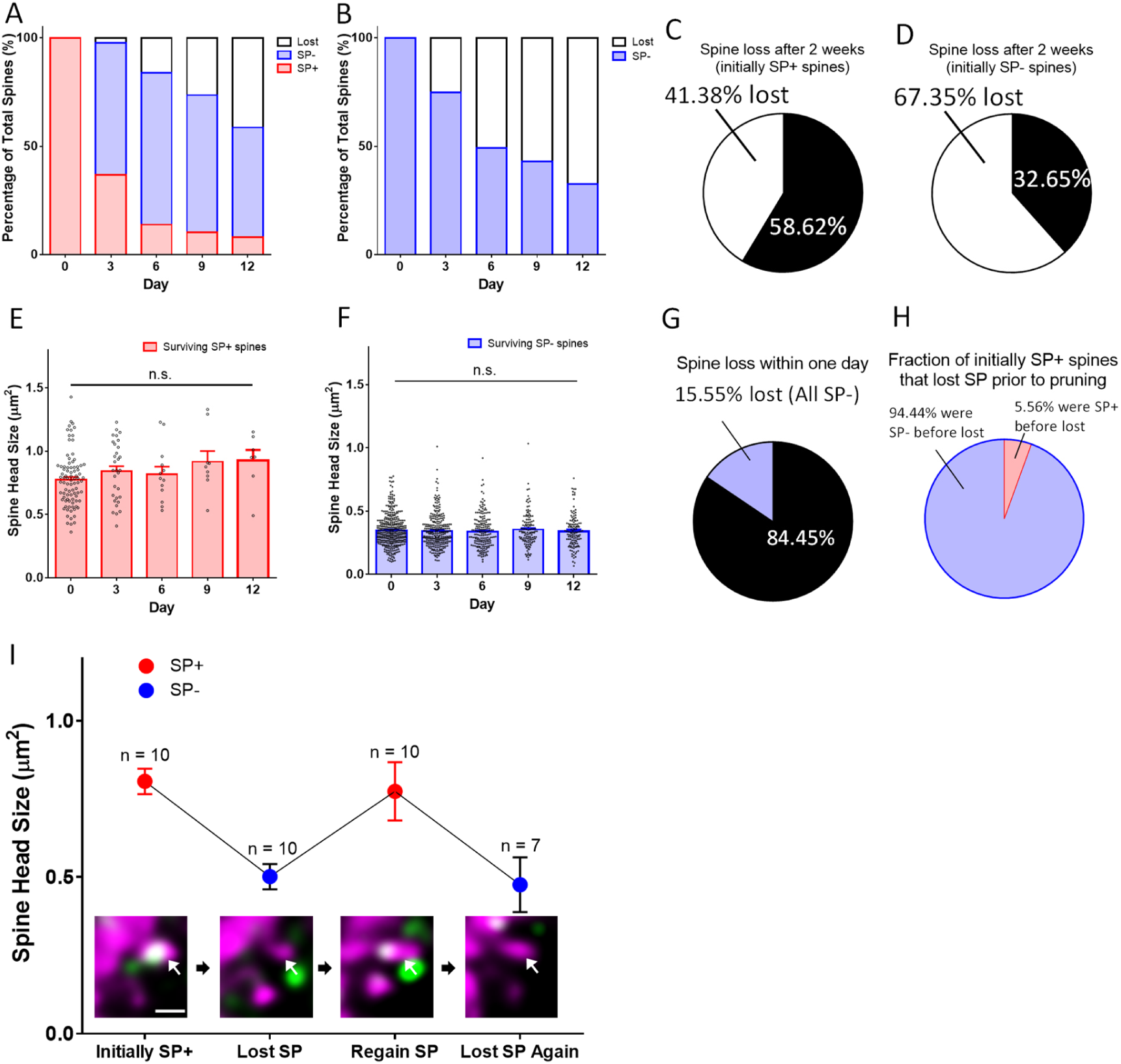
Spines pass through a SP-negative state before pruning. (A, B) The fates of (A) a group of SP+ spines (n = 87) and (B) a group of SP- spines (n = 392) are illustrated. SP+ spines gradually become SP- or disappear. SP- spines disappear with time. (C, D) Fractions of (C) SP+ and (D) SP- spines surviving for two weeks. (E, F) Mean spine head sizes of surviving (E) SP+ and (F) SP- spines. Spine head sizes were not significantly different (SP+ spines, p = 0.115; SP- spines, p = 0.627, Kruskal-Wallis test). (G) Analysis of SP content of all spines lost within one day (days −1 to 0; n = 72; 15.55% of 463 spines). All pruned spines were SP-. (H) Analysis of all SP+ spines that were pruned during the observation period. Most spines (n = 34) went through a SP- state before disappearing, a minority (n = 2) was lost directly between imaging days 0 and 3. (I) A subpopulation of SP+ spines undulated between SP+ and SP- states (n = 10). The gain or loss of SP was accompanied by an increase or decrease of spine head size, respectively. An example for such a case is illustrated (arrow in insets). Scale bar = 1 µm. The following figure supplements are available for Figure 7. **Figure S3 related to Figure 7. Time-lapse 2-photon imaging data of individual SP+ spines.** **Figure S4 related to Figure 7. Time-lapse 2-photon imaging data of individual SP- spines.** **Figure S5 related to Figure 7. Time-lapse 2-photon imaging data of individual spines undulating between SP+ and SP- states.**

In addition, the spine head sizes of the surviving SP+ (Fig. 7E) and SP- (Fig. 7F) spines were analyzed for each time point. Spines that repeatedly changed their SP content were excluded and analyzed separately (see below).

This analysis revealed that in the group of initially SP+ spines the fraction of SP+ spines decreased with time whereas the fraction of SP- spines increased. This was followed by an increase in the fraction of lost spines (Fig. 7A). In the group of the initially SP- spines the fraction of surviving spines decreased much more rapidly (Fig. 7B). At the observation endpoint (day 12), ∼59% of spines which were initially SP+ survived (Fig. 7C), whereas only ∼33% of spines which were initially SP- were still present (Fig 7D). The size of the surviving SP+ (Fig. 7E) and SP- (Fig. 7F) spines did not change significantly during the two-week observation period, in line with our finding that spine head size per se is not a major determinant of spine stability.

We then looked specifically at the spines that were lost, since our analysis had revealed a shift from SP+ spines to SP- spines (Fig. 7A). This shift suggested that SP+ spines go through a SP- state before pruning. Since we followed the fate and the SP content of every spine over the two-week imaging time period (Suppl. Figs. 3, 4 related to Fig. 7), we could test this hypothesis. First, we determined the SP content of the 16% of spines that were lost within one day, i.e. between days −1 and 0. All of these pruned spines were SP-, in line with our hypothesis (Fig. 7G). Next, we predicted that SP+ spines should not be lost without going through a SP-state. For this analysis, we looked at all spines that were initially SP+, i.e. spines SP+ at day 0, and focused on those SP+ spines that were pruned sometime during the observation period. Of these spines ∼94% were SP- prior to their pruning (Fig. 7H). In the rare cases (n=2; ∼6%), in which a SP+ spine seemingly disappeared without passing through a SP- state, the two observation time points were several days apart (e.g., between day 0-3; Suppl. Fig. 3 related to Fig. 7), making it highly likely that we missed a transitory SP- state of these spines prior to their pruning.

### Loss of SP from spines is not sufficient to cause pruning

After finding the above evidence that SP is removed from spines prior to their pruning, we wondered whether removal of SP from a spine is inexorably followed by the loss of this spine. If this were the case all SP+ spines losing their SP cluster should disappear. This hypothesis could be falsified, since we observed several SP+ spines (n=10) undulating between a SP+ and SP- state (Fig. 7I). Thus, SP can be removed from a spine and reintroduced into a spine without the spine being pruned. Of note, the changes in SP content were accompanied by bidirectional changes in head size (Fig. 7I, Suppl. Fig. 5 related to Fig. 7). We also observed SP- spines, which became SP+, lost SP, and became SP+ again at a later stage (n=8; data not shown). Taken together, we conclude from these observations that loss of a SP cluster from a spine is not sufficient to cause pruning of that spine.

### Mathematical simulations of spine stability are compatible with a conditional two-stage decay process

The decay curves of SP- and SP+ spines (Fig. 4, 5) are non-overlapping and mathematically distinct since they cannot be trivially transformed into each other, e.g. using a scaling factor. We first analyzed the survival curve of SP- spines, which we could readily simulate and fit to our data using a one-phase exponential decay function:

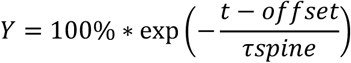

In this equation, the variable Y denotes the percentage of SP- spines surviving up to day t starting with 100% SP- spines present at day zero. τspine denotes the decay time constant of spine loss. The half-life of spines was determined as 0.69*τspine. “offset” denotes the start of the decay. Fitting the data to this model, τspine of SP- spines is ∼9.8 days and the half-life of these spines is ∼6.8 days (Fig. 4). Size matched SP- spines (Fig. 4G, H), SP- spines of different sizes (Fig. 5), and SP-deficient spines (Fig. 6) followed similar one-phase exponential decay functions.

In contrast, the survival curve of SP+ spines did not follow a one-phase exponential decay function. An almost perfect fit could be obtained, however, by using a conditional double phase exponential decay function, identical to the one used to model the conditional two-stage decay of radioisotopes (Bateman, 1910). In this model, a radioisotope undergoes decay and forms an instable intermediary. This intermediary decays again into a stable isotope. The two decay processes have different decay time constants. We now applied this function to the data obtained for spine survival: SP+ spines were considered to lose SP with the time constant τSP. The new SP- spine would then disappear with the time constant τspine, i.e. the same time constant determined for the SP- spines above:

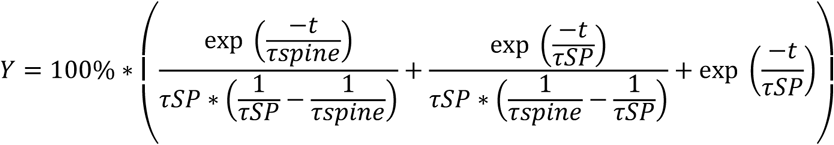

In this equation, the variable Y denotes the percentage of SP- spines surviving up to day t starting with 100% of SP+ spines present at day zero. We fitted the survival data for SP+ spines to this equation and found the time constant τSP of the first decay process (SP+ spines becoming SP-) to be ∼11.1 days and accordingly the half-life of SP in spines to be ∼7.7 days. Thus, after approximately 17.4 days, ∼50% of all initially SP+ spines would have disappeared (Fig. 4). Size matched SP+ spines (Fig. 4G, H) and SP+ spines of different sizes (Fig. 5) followed similar conditional double phase exponential decay functions.

We conclude from this mathematical analysis of our experimental data that the survival curves of SP+ spines are compatible with a conditional two-stage decay process in which SP+ spines first lose their SP cluster (τSP) before being subsequently pruned as SP- spines with the time constant (τspine; same as initially SP- spines; Fig. 8).

**Figure 8.**
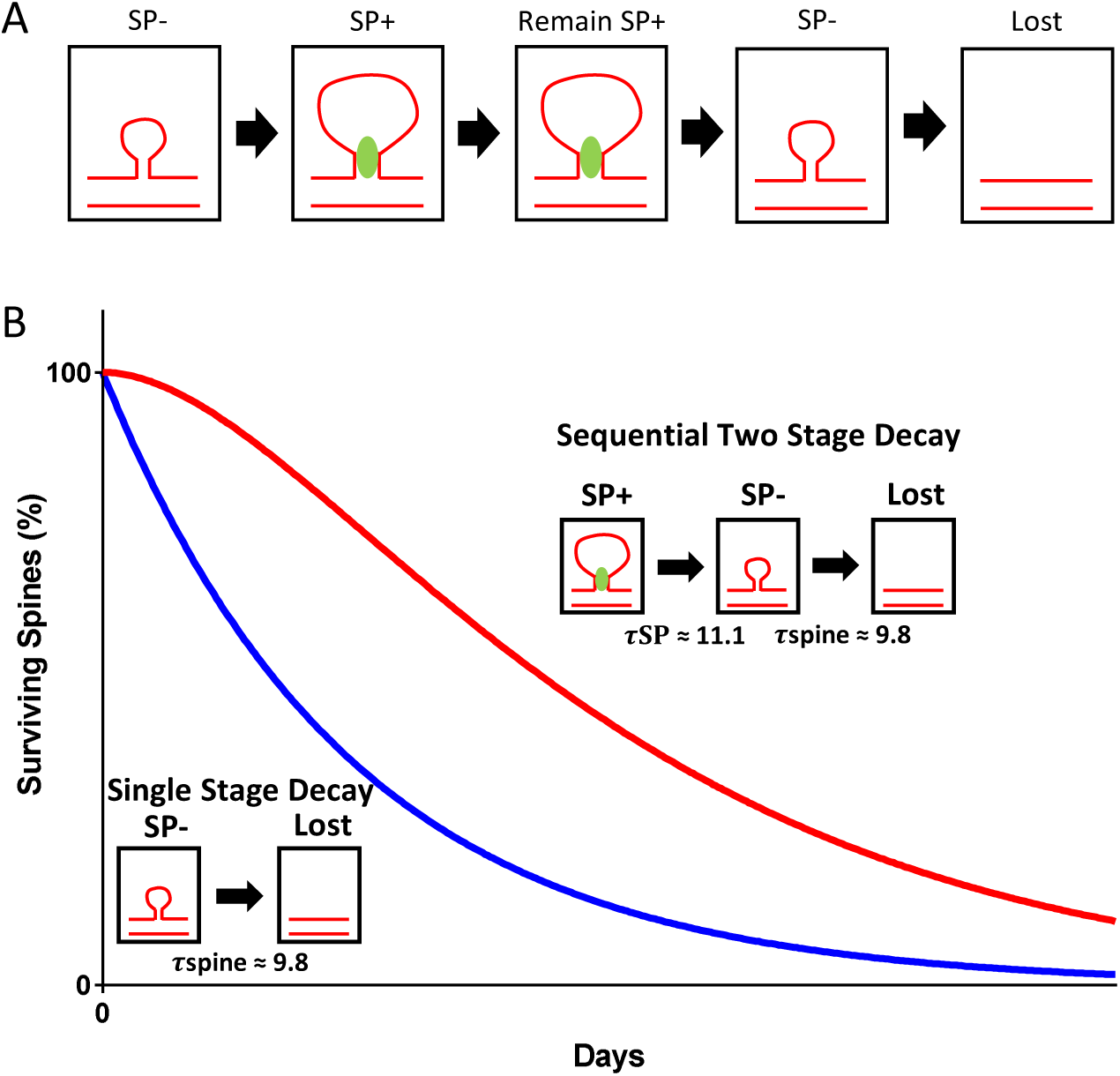
Model illustrating the role of SP in spine head size and spine stability. (A) SP- spines gaining SP show a concomitant increase in spine head size. SP in spines stabilizes spines. Prior to spine loss, SP is removed from spines. Loss of SP from spines is associated with a reduction in spine head size. (B) The data of the present study suggest a conditional two-stage exponential decay model for SP+ spines. The first stage involves SP+ spines losing SP (τSP) and the second stage involves the gradual disappearance of these SP- spines (τspine). For SP- spines, our data suggest they undergo a classical single stage exponential decay model where spines are lost exponentially with time.

## Discussion

Spine geometry and spine stability are structural parameters that have been linked to synaptic strength, network reorganization and memory trace formation (Bourne and Harris, 2007; Kasai et al., 2010; McKinney, 2010; Koleske, 2013; Rogerson et al., 2014; Segal, 2017). A molecule potentially involved in these biological phenomena is the actin-modulating protein SP (Mundel et al., 1997; Deller et al., 2000a; Jedlicka and Deller, 2017). Using a combination of mouse genetics, viral transduction and 2-photon time-lapse imaging, the effects of SP on spine geometry and stability were analyzed for two weeks in an organotypic environment. Our observations can be summarized as follows: (1) SP is primarily present in large spines. (2) SP+ spines are more stable than SP- spines. (3) The effect of SP on spine stability is independent of spine size and is seen in small as well as in large spines. (4) In SP- deficient animals, spines decay like SP- spines. In these mutants, a compensatory increase in spine turnover is observed. (5) Analysis of SP+ spines that were subsequently pruned revealed that these spines went through a SP- state prior to their removal, following a conditional two-stage decay curve. (6) Removal of SP from spines is not sufficient to induce spine pruning.

In sum, our results implicate SP as a major regulator of spine stability. Since SP is primarily present in large spines, its presence explains why these spines have a considerably longer lifetime than small spines, as reported by others (e.g., (Matsuzaki et al., 2004; Kasai et al., 2010; McKinney, 2010).

### Spine head expansion and shrinkage is correlated with SP content and SP cluster size

SP is an actin-modulating protein (Mundel et al., 1997; Asanuma et al., 2006; Okubo-Suzuki et al., 2008). Since geometry and dynamics of spines depend on F-actin-assembly and disassembly (Fischer et al., 1998; Matus, 2000), a role for SP in spine motility appeared to be likely (Deller et al., 2000a). Indeed, after investigations in SP-deficient mice showed that SP does not influence spine density or spine length (Deller et al., 2003), a robust correlation between SP and spine head size was reported in dissociated hippocampal neurons transfected with SP (Okubo-Suzuki et al., 2008; Vlachos et al., 2009) and in acute hippocampal brain slices (Zhang et al., 2013). In our organotypic tissue culture preparations we have confirmed and extended these findings and observed a strong correlation between SP content and spine head size as well as between SP cluster size and spine head size. Furthermore, the mean cross-sectional area of SP+ spines was found to be almost twice as large as the mean cross-sectional area of SP- spines. By following single SP+ spines over several days, we could also detect a group of SP-containing spines that lost and subsequently regained SP. The size of the spine head shrank in spines losing SP and increased in spines regaining SP. Thus, SP content and cluster size correlated with changes in spine head size in both directions. However, SP itself does not seem to cause spine head-size expansion, since short-term imaging revealed that spines growing bigger heads first expand their heads and subsequently insert SP (Okubo-Suzuki et al., 2008; Konietzny et al., 2019), maintaining the expanded spine head (Okubo-Suzuki et al., 2008).

### The presence of SP rather than head size determines spine stability

Spines differ with regard to their lifetime and stability. Although spines of all sizes can persist for long time periods, spines with a large spine head are believed to be more stable than spines with only a small head (Matsuzaki et al., 2004; Kasai et al., 2010). Alternatively, if classifications of spine shape are used to categorize spines (Bourne and Harris, 2008), mushroom spines were reported to be more stable than stubby or thin spines (Kasai et al., 2003; Bourne and Harris, 2007; Koleske, 2013), even following denervation (Caceres and Steward, 1983). Since mushroom spines often have large spine heads, a considerable overlap exists between “spines with a large spine head” and “mushroom spines”, explaining the similarity of the results. Regardless of these different classifications, however, spine size has been considered a major determinant of spine stability (Kasai et al., 2010; McKinney, 2010).

Here we have investigated the molecular basis for this increased spine stability and have tested the possibility that the actin-modulating protein SP (Mundel, 1998; Deller et al., 2003) could play a role. Since spine stability depends on the stability of the spine actin cytoskeleton (Fischer et al., 1998; Matus, 2000), actin-modulating proteins such as SP are candidate regulators for spine motility and stability (Deller et al., 2000a; Bourne and Harris, 2008). Our data revealed that SP is indeed a major regulator of spine stability. Furthermore, our data also suggest that it is primarily the content of SP within a spine that imparts stability to a given spine rather than the size of the spine per se. This second conclusion is supported by the following: (1) Small spines (0.3-0.55 µm^2^ head area) containing SP have a long lifetime and a median survival of 12.8 days. Large spines (>0.8 µm^2^ head area) without SP have a median survival time of 7.7 days. Thus, small SP+ spines are more stable than large SP- spines. (2) Comparison of the median survival times of SP- spines of all sizes (i.e., very small, small, medium and large) revealed that their median survival time was similar and much shorter than the median survival time of any of the SP+ spine groups. Finally, (3) all spines, including the large spines, of SP-deficient animals decayed like SP- spines and had short median survival times. Although we cannot fully exclude a small effect of spine size on spine survival, e.g. large spines may need time to shrink prior to their removal (see next paragraph), our data clearly show that the presence of SP within a spine is a more reliable indicator of its stability than its morphology alone, i.e. the size of its spine head.

### SP+ spines that are pruned pass through a SP- state and shrink in size

Two weeks of time-lapse imaging of identified SP+ spines made it possible to study their fate: Single SP+ spines surviving for the entire time period, SP+ spines losing and regaining SP, as well as SP+ spines that were pruned could be identified and analyzed (Suppl. Figs. 3, 4, 5 related to Fig. 7). From these multidimensional dynamic time-lapse data, we could draw additional conclusions: First, from the fact that some spines lose and regain SP with time, we concluded that loss of SP from a spine is not by itself sufficient to tag a spine for pruning. Rather, such spines seem to re-enter the large pool of SP- spines from which new SP+ spines can be recruited. Second, by following SP+ spines that were pruned during the observation period, we could show that in almost all cases, these spines went through a SP- state before they disappeared. In the two cases in which SP+ spines were pruned without an intermediate state, the two observation time points were several days apart (e.g. three days between day 0 and day 3), making it highly likely that the intermediate SP- state was missed. Thus, we conclude that removal of the spine stabilizing protein SP is a necessary step before the actin-cytoskeleton of a spine can be degraded and the spine can be pruned. Third, in line with the fact that SP is associated with large spines, we observed that spines losing SP also shrank in size (Suppl Figs. 3-5, related to Fig 7). In the case of spines changing their SP content (Suppl. Fig. 5, related to Fig. 7) this also resulted in several corresponding head size changes. In agreement with these observations, we found that SP+ spines designated to be pruned follow a conditional two-stage decay function, similar to the one described for radioactive isotopes in Physics (Bateman, 1910): The first stage is the “decay” of SP+ spines into a SP- state. The second stage of the decay is the pruning of these SP- spines, which is as fast as for other spines that were primarily devoid of SP (Fig. 8).

### SP-deficient neurons homeostatically compensate for the loss of SP with increased spine formation

The role of SP in the regulation of spine stability is further supported by data from SP-deficient mice. Granule cells of these mice exhibited spines that decayed like SP- spines. In addition, SP-deficient granule cells compensated for the reduced stability of spines by increasing their spine formation and thereby their spine turnover ratio. This finding suggests that SP-deficient neurons homeostatically compensate for the loss of stability by upregulating their spine formation. This may explain why loss or overexpression of SP does not seem to affect spine densities (Deller et al., 2003; Okubo-Suzuki et al., 2008; Vlachos et al., 2009). Whereas spine density appears to be regulated by other molecules, including among others hormones (Woolley, 2000; Segal and Murphy, 2001; Fester et al., 2012; Luine and Frankfurt, 2012), actin-modulating proteins (Yamazaki et al., 2014), Fragile X mental retardation protein (Irwin et al., 2000; Bagni and Zukin, 2019), non-coding RNAs (Briz et al., 2017), neurotrophic factors (Murphy et al., 1998) and adhesion molecules (Segura et al., 2007; Keeler et al., 2015; Muller et al., 2017), SP appears to be a major regulator of spine stability.

### SP marks a subpopulation of stable spines - implications for synaptic plasticity and long-term memory

The present study showed that SP is found in a subpopulation of granule cell spines that are highly stable. The stabilizing effect of SP is most likely caused by its actin-modulating function: In hippocampal neurons, it has been shown to protect F-actin from disruption (Okubo-Suzuki et al., 2008; Wang et al., 2016) and in kidney it has been demonstrated to regulate actin organization by protecting RhoA from Smurf1-mediated ubiquitination and proteasomal degradation by competing with Smurf1 for RhoA binding (Asanuma et al., 2006). Since the Rho family of small GTPases regulate the maintenance of spines and spine formation (Tashiro and Yuste, 2008), a similar role for SP in the CNS has been suggested (Asanuma et al., 2006; Zhang et al., 2013). However, the kidney form of SP differs from the neuronal form (Asanuma et al., 2006), and thus this attractive hypothesis still awaits confirmation.

SP also plays an important role in Hebbian- (Yamazaki et al., 2001; Deller et al., 2003; Okubo-Suzuki et al., 2008; Holbro et al., 2009; Jedlicka et al., 2009; Vlachos et al., 2009; Zhang et al., 2013; Jedlicka and Deller, 2017) and homeostatic (Vlachos et al., 2013) forms of synaptic plasticity. These effects on plasticity may not only depend on the actin-modulating properties of SP but also on the spine apparatus organelle, which requires SP for its formation (Deller et al., 2003). The spine apparatus is part of the effector machinery needed for changing synaptic strength at excitatory postsynapses (Jedlicka and Deller, 2017). It acts as a calcium source and sink within spines (Vlachos et al., 2009; Korkotian and Segal, 2011; Korkotian et al., 2014) and is required for AMPA-R trafficking to the postsynapse (Vlachos et al., 2009). Thus, SP appears to be involved in at least four biological phenomena linked to changes in synaptic strength and in the maintenance of such a change, i.e. prolonged spine head expansion (Okubo-Suzuki et al., 2008; Vlachos et al., 2009; Zhang et al., 2013), spine stabilization (this study), calcium release from intracellular stores (Vlachos et al., 2009; Korkotian and Segal, 2011; Korkotian et al., 2014) and AMPA-R trafficking (Vlachos et al., 2009), demonstrating its important role as a plasticity-related protein. Conversely, the presence of SP in spines identifies spines that are stable, large and strong.

At the network level, strong and stable synapses are considered the backbone of long-term memory (Kasai et al., 2010; Koleske, 2013; Rogerson et al., 2014; Segal, 2017). Neurons connected by such synapses form ensembles of cells that are in turn essential for memory engrams (Yokose et al., 2017). Indeed, as has been shown recently, the sharing of engram cells can link memories, whereas synapse-specific plasticity guarantees the identity and storage of individual memories (Abdou et al., 2018). In the present study, we report that the stability of spines critically depends on the presence of the actin-modulating protein SP within a spine. This makes it attractive to speculate that synapses requiring a high degree of stability, such as those synapses binding excitatory neurons together into functional ensembles, require SP as their stabilizer. Conversely, SP may be an attractive marker for such synapses, as it allows for their identification in vitro and possibly also in vivo. Using optogenetic approaches, such as the one employed by Abdou and co-workers (Abdou et al., 2018), the role of SP+ spines in memory formation can now be addressed.

## Acknowledgement(s)

We thank Charlotte Nolte-Uhl and Anke Biczysko for technical support and Dr. Stephan W. Schwarzacher for critical discussion of our data. The authors (D.D.T., A.D., T.D.) dedicate this paper to the memory of the late Michael Frotscher, M.D., who was a wonderful mentor and an enthusiastic scientist and who was involved in many pioneering studies on the localization and the role of Synaptopodin in the CNS.

## Supplementary Figures and Captions

**Figure S1 related to Figure 2.**
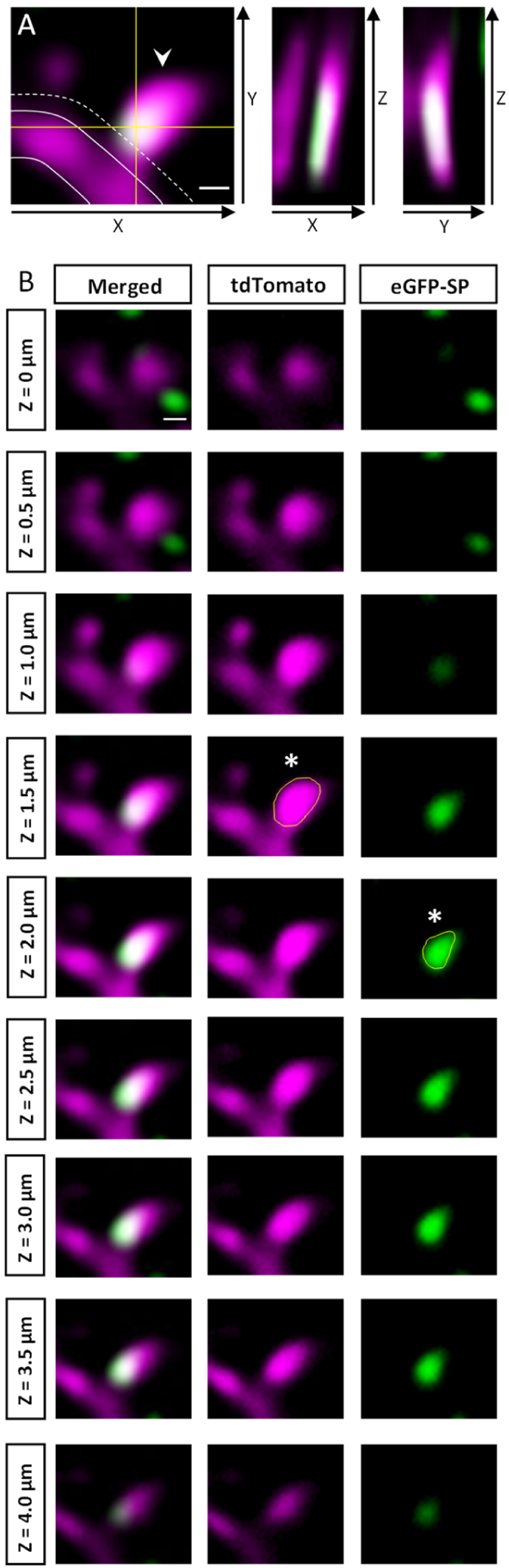
Measurement of dendritic spine head size and SP cluster size. (A) Only protrusions exceeding the dendritic shaft (outlined) laterally in the x-y directions for at least 0.35 µm (dotted line) were included in the analysis. Spines were considered SP+ (arrowhead) if the SP cluster overlapped with the spine head and/or neck in both the x-z and y-z directions when scrolling through the z-stack. Scale bar = 0.5 µm. (B) A series of x-y planes taken at consecutive z-levels is illustrated. Spine head size and SP cluster size are measured by choosing the planes with the maximal cross-sectional area of the spine head and SP cluster when going through the z-stack. x-y planes with the largest cross-sectional area of spine head (middle column, yellow outline of spine head, asterisk) and SP cluster (right column, yellow outline of SP-cluster, asterisk) are highlighted. Scale bar = 0.5 µm.

**Figure S2 related to Figure 6.**
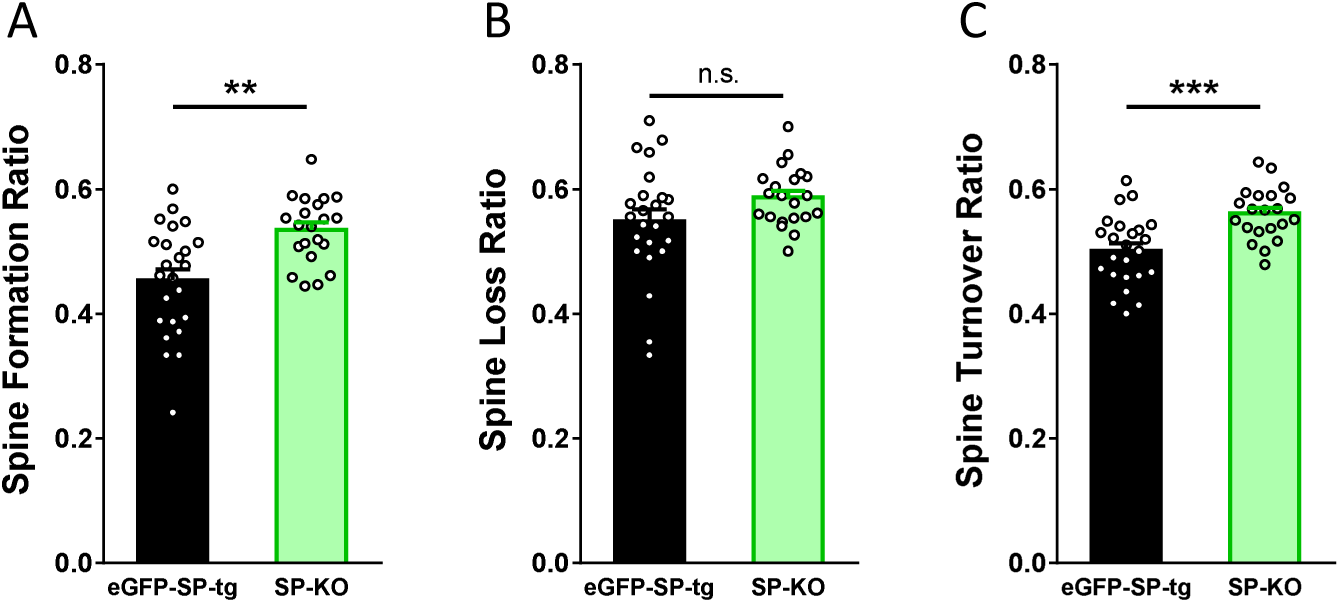
SP-KO mice compensate for the loss of SP with an increased spine formation and an increased turnover ratio. (A) Spine formation was significantly higher in SP-KO dendrites as compared to eGFP-SP-tg dendrites. **p = 0.0011, Mann-Whitney U-test. Number of dendritic segments: eGFP-SP-tg, n = 24; SP-KO, n = 21. (B) Spine loss was not significantly different between SP-KO and eGFP-SP-tg dendrites. p = 0.0868, Mann-Whitney U-test. Number of dendritic segments: eGFP-SP-tg, n = 24; SP-KO, n = 21. (C) Spine turnover ratio was significantly higher in SP-KO dendrites as compared to eGFP-SP-tg dendrites over the two-week observation period. ***p = 0.0003, Mann-Whitney U-test. Number of dendritic segments: eGFP-SP-tg, n = 24; SP-KO, n = 21.

**Figure S3 related to Figure 7.**
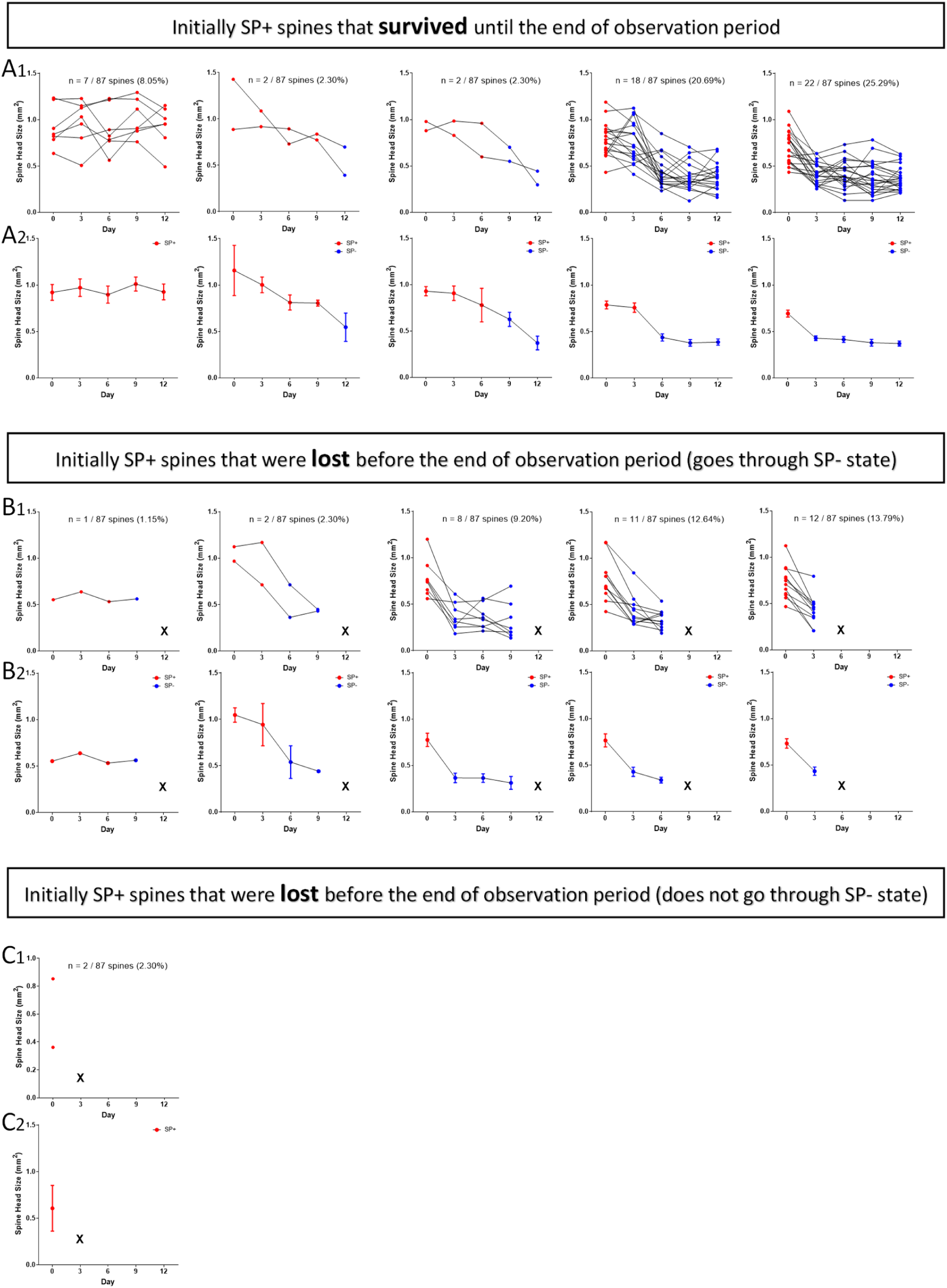
Time-lapse 2-photon imaging data of individual SP+ spines. SP+ spines (SP+ on day 0) were subdivided into three categories: (A) spines that survive until the end of the observation period, (B) spines that are pruned and go through a SP- state, (C) spines that are pruned without going through a SP- state before pruning. The fractions, percentages and SP-content (SP+, red; SP-, blue) of spines in the different categories are indicated. Upper panels (A1 through C1) show individual spines, lower panels (A2 through C2) show summary diagrams (mean ± SEM) of data illustrated in A1 through C1. Spines undulating between SP- and SP+ states were excluded and analyzed separately (c.f. Suppl. Fig. 5). (A1, A2) Initially SP+ spines surviving until the end of the observation period (n = 51). Spines are sub-grouped based on the day SP is lost from the spine. Note the reduction of spine head size after loss of SP. (B1, B2) Initially SP+ spines that are pruned and go through a SP- state prior to pruning (n = 34). X = lost. Spines are sub-grouped based on the day the spine is pruned. (C1, 2) Initially SP+ spines that are lost directly (n = 2; 5.56% of SP+ spines), i.e. without going through a SP- state. These rare spines disappear sometime between imaging days 0 and 3.

**Figure S4 related to Figure 7.**
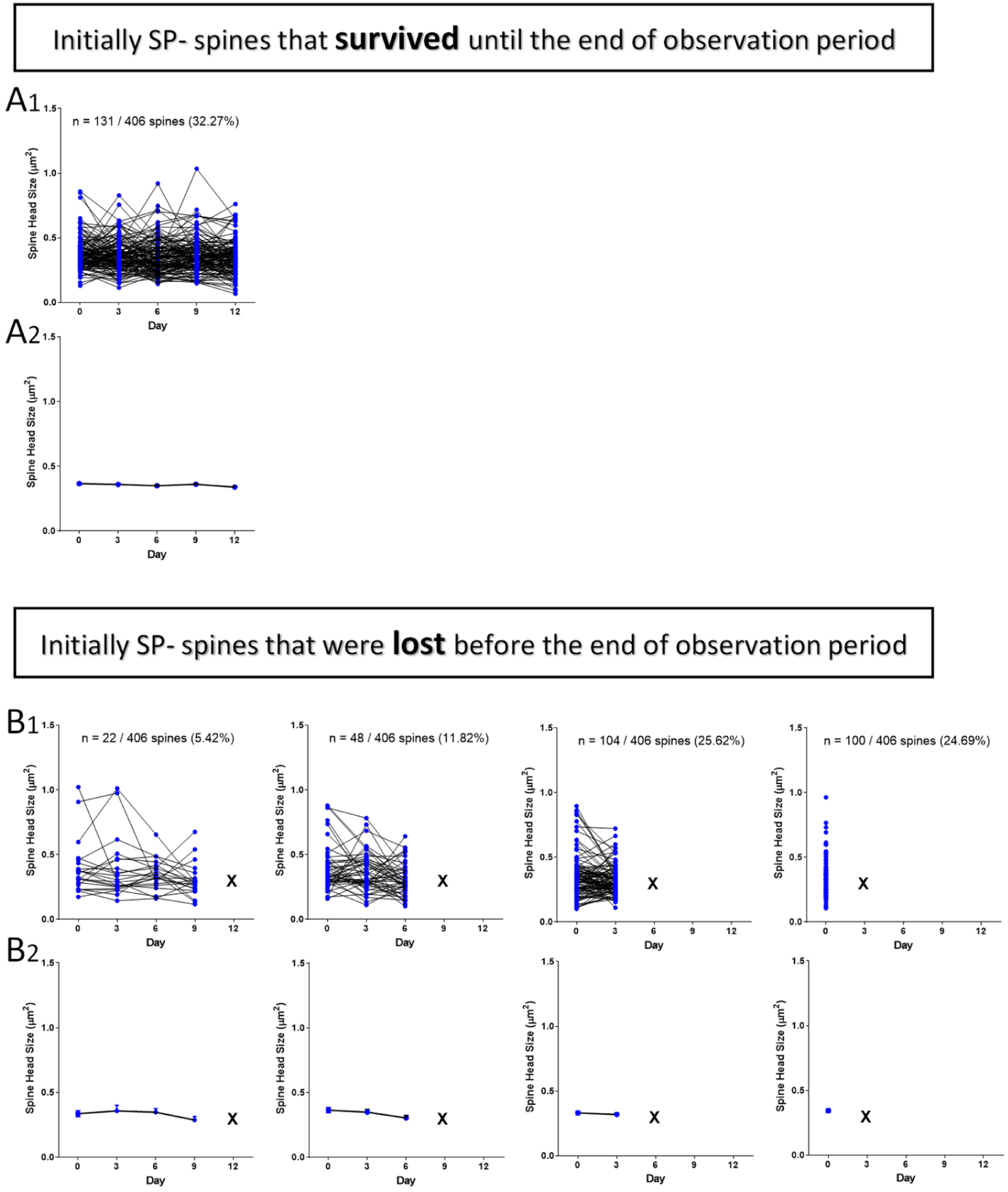
Time-lapse 2-photon imaging data of individual SP- spines. SP- spines (SP- on day 0) are subdivided into two categories: (A) spines that survive until the end of the observation period (n = 131), (B) spines that are lost (n = 275). The fractions, percentages and SP-content (SP-, blue) of spines in the different sub-categories are shown. Upper panels (A1 and B1) show individual spines, lower panels (A2 and B2) show summary diagrams (mean ± SEM) of data shown in A1 and B1. Spines undulating between SP+ and SP- states are excluded and analyzed separately (c.f. Suppl. Fig. 5).

**Figure S5 related to Figure 7.**
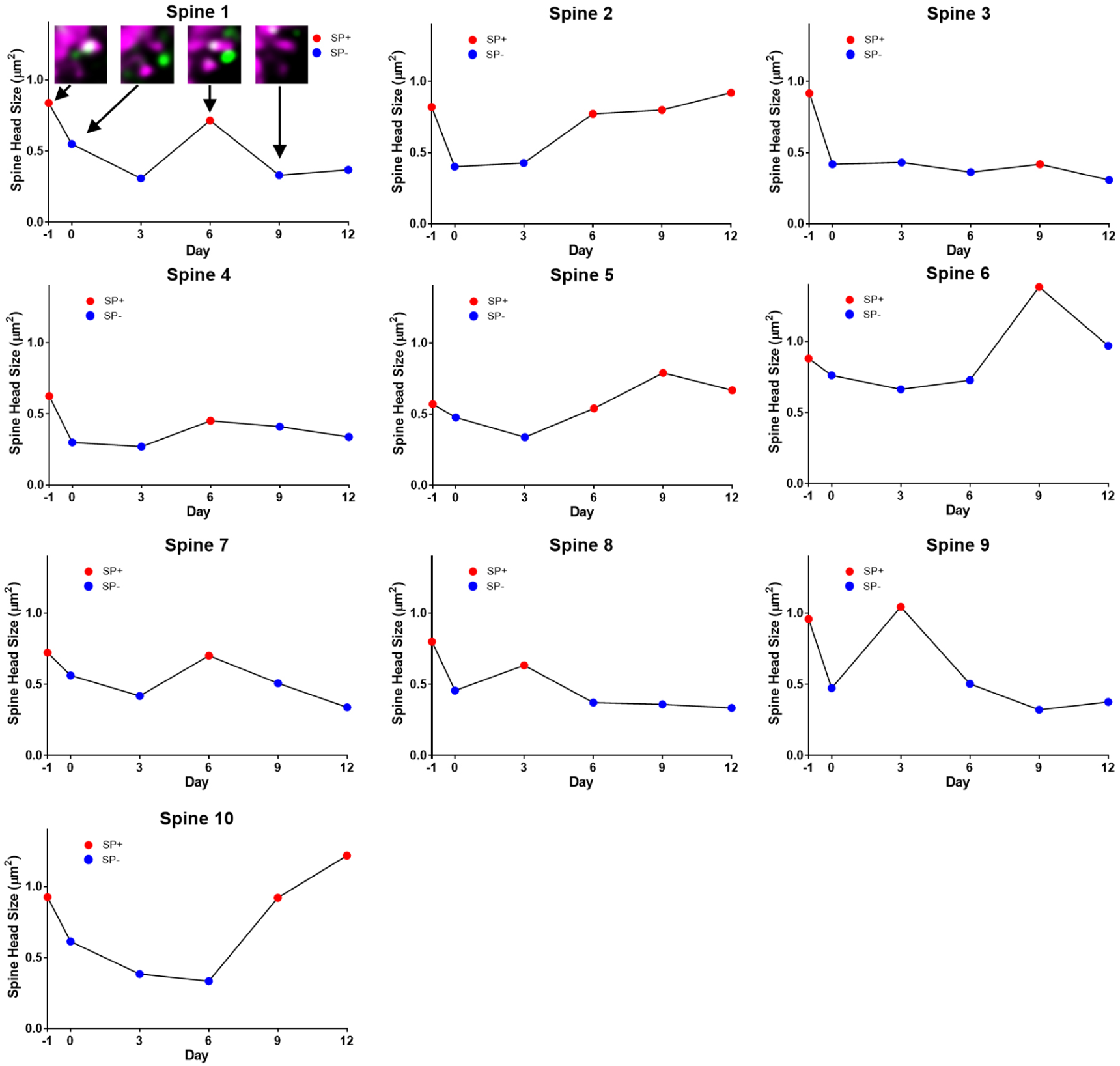
Time-lapse 2-photon imaging data of individual spines undulating between SP+ and SP- states. Some SP+ spines loose SP and subsequently regain SP. This demonstrates that loss of SP from spines is not sufficient for spine pruning. Note corresponding changes in SP-content and in spine head size. The example of an undulating spine illustrated in Fig. 7I (lower part) is shown in this figure with all imaging time points (Spine 1).

